# Integrative immune subtyping of HNSCC reveals clinically relevant phenotypes and treatment-associated transitions

**DOI:** 10.1101/2025.07.31.667850

**Authors:** Indu Khatri, Terezinha de Souza, Saskia D van Asten, Merzu Belete, Felipe Vieira Braga, Merel Jongmans, Sri Sridhar, Brandon W. Higgs, Iris Kolder

**Author notes:** These authors contributed equally (Terezinha de Souza, Saskia D van Asten). Correspondence to: Indu Khatri and Iris Kolder, Genmab B.V., Uppsalalaan 15, 3584 CT, Utrecht, The Netherlands.

## Abstract

Head and neck squamous cell carcinoma (HNSCC) exhibits profound heterogeneity in clinical presentation, treatment response, and immune landscape. While prior classification systems have identified molecular and immune subtypes in this disease, their applicability to real-world clinical settings remains restricted to small, homogeneous cohorts and limited by lack of multimodal data integration and interpretation.

We performed integrated multi-omics analysis including transcriptomic, genomic (copy number, single-nucleotide variants) on 1,149 tumors from 1,102 HNSCC patients across treatment settings. Using the Similarity Network Fusion (SNF) algorithm, we defined immune subtype clusters (ISCs) based on the full immune gene landscape. These clusters were characterized using mutational, transcriptional, and immune cell enrichment analyses, and mapped to hypoxia and traditional subtypes. Associations with clinical outcomes, including progression-free survival, were evaluated across first-line and post-metastatic treatment settings.

Four distinct immune subtype clusters (ISC1–ISC4) were identified: ISC1: immune-cold and EMT-enriched; ISC2: immune activated; ISC3: mixed immune-regulatory and stromal-enriched phenotype; and ISC4: immunosuppressed. Distinct treatment response patterns were observed across subtypes in subjects treated with checkpoint inhibitors, chemotherapy, and combination regimens. 44 Patients with matched pre/post treatment tumors revealed treatment-associated transitions between immune subtypes: checkpoint inhibitor treatment enriched for immune activation, while chemotherapy treatment enriched for immunosuppressive signaling pathways.

This study provides a clinically relevant immune subtyping framework for HNSCC based on real-world, multi-omics data. These subtypes reflect dynamic tumor–immune states and associated with treatment response and survival, supporting their use in guiding immune-based therapy in HNSCC.

## Introduction

Head and neck squamous cell carcinoma (HNSCC) originates from the mucosal epithelium of the pharynx, larynx, oral cavity, or sinonasal tract (1). Globally, HNSCC is the seventh most common cause of cancer and accounts for ∼325,000 deaths annually (2). Known risk factors include tobacco and alcohol use, as well as human papillomavirus (HPV) infection (1,3). Despite advancements in treatment options such as surgery, chemotherapy, and radiotherapy, clinical outcomes remain poor for many patients, with the 5-year survival rate averaging 50% globally (2).

Targeted therapies, including EGFR inhibitors (4,5) and immune checkpoint inhibitors (CPI) (6,7), have been increasingly utilized in HNSCC, yet their effectiveness is limited to specific patient subgroups, with many patients failing to respond. The challenges in treatment response are largely attributed to the substantial heterogeneity of HNSCC at the etiological, phenotypic, and molecular levels (8,9). This underscores the urgent need for a deeper understanding of immune heterogeneity in HNSCC, particularly in real-world datasets where patient diversity better reflects clinical practice.

Efforts have been made to classify HNSCC into molecular subtypes, with studies such as The Cancer Genome Atlas (TCGA) identifying four distinct subtypes: basal, mesenchymal, atypical, and classical (10). These classifications have significantly advanced our understanding of HNSCC biology and provided a framework for studying tumor microenvironment (TME) and treatment response. Additionally, immune profiling studies have sought to categorize HNSCC into immune subtypes, typically identifying three major clusters—immune-inflamed, immune-excluded, and immune-desert phenotypes—each with distinct implications for immunotherapy responsiveness (11,12).

In addition to immune heterogeneity, hypoxia plays a crucial role in shaping the TME and modulating immune responses in HNSCC. Hypoxic tumors commonly exhibit immunosuppressive features, including reduced cytotoxic T-cell infiltration, increased recruitment of regulatory T cells and myeloid-derived suppressor cells (MDSCs), and upregulation of immune checkpoint molecules, contributing to resistance against immunotherapy (13). Hypoxia subtypes, defined through transcriptomic profiling (14), have been linked to tumor aggressiveness and poorer outcomes While these classifications have been instrumental in defining molecular and immune landscapes, most studies have been derived from controlled datasets with relatively homogeneous patient populations, limiting their applicability to real-world settings (15). Moreover, immune subtyping has largely relied on transcriptomic data, often focusing on predefined immune signatures rather than using a broader, integrative approach.

To build upon these foundational studies to a broader clinical context. We applied Similarity Network Fusion (SNF) (16) to transcriptomic, copy number, and variant data from real-world patient samples. Unlike prior studies that required feature space reduction, our approach preserved the full immune gene landscape to identify ISCs. These subtypes were further assessed against molecularly defined hypoxia subtypes to characterize the interplay between hypoxia-driven adaptations and immune states, providing a more comprehensive view of immune heterogeneity in HNSCC. To explore the evolution of immune subtypes over time, we also analyzed paired biopsies from longitudinal patient samples and integrated single-cell data to evaluate immune transitions during tumor progression. This classification system complements earlier efforts and enables more granular stratification to guide immunotherapy and targeted treatment decisions.

## Material and Methods

### Cohort and metadata description

We analyzed de-identified patient records with a primary diagnosis of HNSCC in a cohort of 1,149 real-world (RW) tumor biopsies from 1,102 patients (**Table 1**) where 42 patients had more than one biopsy available. This included 20 patients with paired pre– and post-treatment biopsies, 15 with paired post-treatment biopsies, and seven with both biopsies collected pre-treatment. Additionally, one patient had three post-treatment biopsies, and another had four biopsies collected from treatment-naïve sites. Samples were categorized into three Categories: *Naïve (naïve HNSCC biopsies)*, *Indication (Naïve + treated HNSCC biopsies)* and *All Samples (full cohort)* to minimize confounding by sample origin. Sample-level clinical metadata were assigned to biopsies, however when the patient-level longitudinal annotations were available the closest available event was assigned to the biopsy.

**Table 1:**
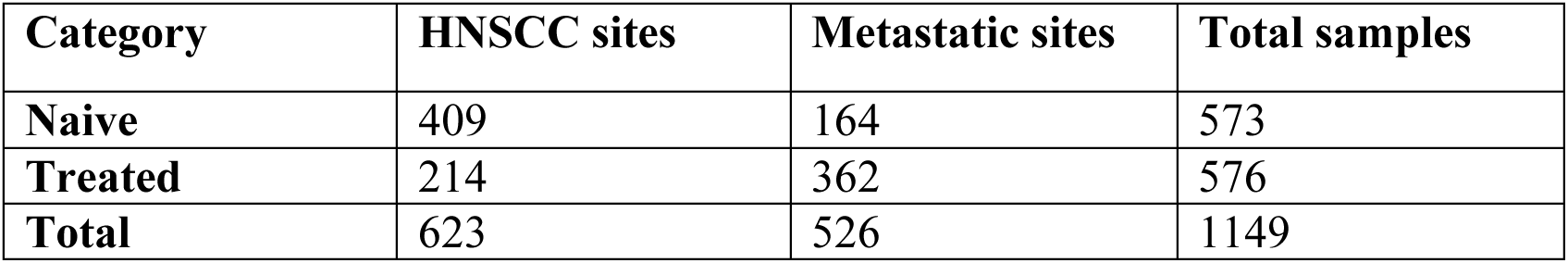
Distribution of biopsies in RWE data in different categories.

### Preprocessing of Sequencing data

#### RNA-Sequencing data

RNA-Seq raw sequencing data were processed using Kallisto to quantify transcript abundances (17). Transcript counts were filtered with at least 10 counts in 5% of samples. A variance stabilized transformation was applied using DESeq2 (18). Principal component analysis was performed to assess batch effects related to assay version, which were subsequently removed using LIMMA’s removeBatchEffect function via linear modeling (19).

#### DNA-Sequencing data

Somatic NGS testing was conducted on all patients using a commercially available gene panel covering 595 to 648 genes (20,21). This panel detected somatic mutations including single nucleotide variants (SNV), insertions and deletions (Indels), copy number variations (CNVs), structural variations, microsatellite instability (MSI), and tumor mutational burden (TMB). Details regarding sample preparation and bioinformatic analyses have been previously published (20,21).

SNVs. short indels and rearrangements were transformed into a binary matrix for analysis, whereas CNV data was retained in its original form. CNV values that were not reported were assumed to represent diploid status (copy number = 2).

#### PD-L1 status

PD-L1 status was available for 474 patients through clinical testing using the 22C3 anti-PD-L1 antibody (Agilent) (22). The Combined Positive Score (CPS) was calculated as the ratio of PD-L1-positive tumor cells, lymphocytes, and macrophages to the total number of viable tumor cells, multiplied by 100. Samples are categorized as PD-L1 negative (CPS < 1%), PD-L1 low (1%–49%), or PD-L1 high (≥50%).

### Assigning HPV status to additional patients

HPV status was available for 143 patients. To infer HPV status for additional patients, a machine learning (ML) model was developed using P16 as a surrogate marker for HPV status; however, to expand the feature space for the ML model, the STRING database (string-db.org) was used to identify functional partners of P16. This search identified 10 associated genes: CDK4, MDM2, CDK6, TP53, NPM1, CCND1, MYC, CDK2, KRAS, and CCND2 which were used as the feature set for model training (**Supplementary Figure S1)**.

A dataset of 152(143 patients) samples with known HPV status and gene expression, a 70/30% train-test split was used. A 5-fold cross-validated random forest (RF) model was trained on the training data to assess gene feature importance. The five most predictive genes (CDKN2A, CDK2, CCND1, CDK6 and MDM2) were selected for training a Support Vector Machine (SVM) model. Thresholds for high-confidence predictions were determined using mean probabilities from true positive and true negative samples in the test set.

Applying the trained SVM model, high-confidence HPV status was assigned to 489 additional patients (505 samples): 284 patients (295 samples) as HPV-negative and 205 patients (210 samples) as HPV-480 patients (492 samples) could not be assigned an HPV status with high confidence.

### Identifying ISCs in the HNSCC patients

Batch corrected RNA-seq counts (**Supplementary Figure S2**), copy number variations, and variant data representing immune-system related genes were used to identify ISCs using SNF, an unsupervised method that effectively integrates multi-omics data without requiring feature space reduction.

A range of hyperparameters were optimized in SNF to explore optimal solutions. The SNF process was configured with 20 clustering solutions, allowing for robust exploration of clustering outcomes. The similarity network construction was optimized by varying key parameters i.e.

1. *Alpha values* were set to a range of 0.3 to 0.7 in increments of 0.1 to control the balance between within– and cross-network similarity.
2. *k-values* (number of nearest neighbors) were explored from 10 to 30 in steps of 5 to fine-tune local neighborhood selection.
3. *t-values* (number of iterations) were varied from 10 to 30 in steps of 5 to optimize information propagation.

To ensure convergence and minimize instability, the analysis allowed up to 100 retry attempts for network fusion. A two-step SNF scheme was used, while no dropout distribution adjustments were applied due to the absence of missing data.

A total of 20 clustering solutions were initially evaluated, and four clusters were selected as the optimal number for HNSCC, based on consistent findings across three sample categories: *Naïve* (ISC-N), *Indication-specific* (ISC-NT), and *All samples* (ISC). After obtaining the clustering solutions, the Adjusted Rand Index (ARI) was computed to evaluate the agreement between the clustering results, ensuring the consistency and validity of sample assignments across different clustering solutions. Additionally, all solutions were manually assessed to confirm that the clusters represented meaningful groupings and did not consist of clusters with fewer than 10 samples, which would indicate poor clustering resolution.

#### 1. Identifying immune subtypes using SNF

RNA, CNV, and variant data were subsetted to include a total 20,457 immune-related genes (includes pseudogenes, LncRNA and miRNA) identified from ImmuneSigDB (23), which consists of 4,872 gene sets derived from the C7: Immunologic Signature Gene Sets and the Reactome’s immune system pathway R-HSA-168256 (24). Samples were categorized into three groups: 1) *Naïve Category* – Naïve (pre-treatment) samples from head and neck sites. 2) *Indication Category* – Both naïve and treated samples from head and neck sites. 3) *All Samples Category* – All naïve and treated samples from both indication and metastatic sites. These groups are inclusive e.g. *Indication* Category comprises of *Naïve Category* and *All Samples Category* comprises of both *Naïve and Indication category*.

For the *Naïve Category*, SNF clustering consistently returned four clusters. In the *Indication Category*, SNF produced either four or seven clusters; however, the seven-cluster solution included three clusters with fewer than 10 samples, making it unsuitable for analysis. Similarly, in the *All Samples Category*, SNF identified four stable clusters.

#### 2. Identifying traditional and hypoxia subtypes using SNF

RNA data were subsetted to include 200 hypoxia-related genes from the MSigDB HALLMARK_HYPOXIA (25) gene set. Samples in the All Samples Category were analyzed using Similarity Network Fusion (SNF) (16) to identify clusters, and solutions with two clusters were retained, as previously shown. Genes known to be upregulated in hypoxia-high clusters (CA9, CASP14, LOX, GLUT3, SERPINE1, AREG, EREG, CCNB1, KIF14) were used to determine whether a cluster exhibited high or low hypoxia.

To identify the four traditional subtypes i.e. Classical, Atypical, Basal, and Mesenchymal, RNA data were subset to include the 2,500 most variable genes for samples in the Indication Category. Genes known to be relevant for traditional subtypes were used to annotate the respective clusters. Similarity Network Fusion (SNF) initially produced three clusters across all solutions. To achieve four clusters, Non-Negative Matrix Factorization (NMF) was applied, and both clustering solutions were evaluated.

### Molecular and Cellular Profiling of Immune Subtype Clusters

To characterize the molecular and cellular features of ISCs, we applied a multi-layered analytical framework incorporating mutational profiling, copy number variation (CNV), gene expression, pathway and cell type enrichment.

#### Identifying enriched mutations and pathways associated with lost or amplified CNVs in the immune subtype clusters (ISCs)

Single nucleotide variants and short insertions or deletions were filtered to include only those classified as pathogenic or likely pathogenic based on functional annotations. Variant types retained for analysis included frameshift mutations, missense mutations, stop-gained alterations, and other functionally consequential coding events. A binary mutation matrix was constructed by collapsing gene symbols and nucleotide-level changes into unique variant identifiers. Enrichment of specific mutations across defined immune and hypoxia subtypes was assessed using Fisher’s exact test.

CNV data were processed to identify genomic regions exhibiting significant copy number gains or losses. Statistical enrichment of CNVs across ISCs was evaluated using Fisher’s exact test, with p-values adjusted for multiple hypothesis testing via the Bonferroni method. Genes located within significantly amplified or deleted genomic regions were subjected to pathway enrichment analysis using the enrichR package, referencing the Reactome Pathways 2024 database. Enrichment analyses were stratified by immune subtype classification (ISC1–ISC4) and CNV category (gain or loss). The most significantly enriched pathways were prioritized based on adjusted p-values and gene set overlap ratios.

#### Identifying genes associated to the ISCs

Differential expression analysis was performed using normalized gene expression data with a one-vs-rest approach for each ISC, using the *limma* package. Genes with a p-value < 0.05 were considered specific to the corresponding subtype.

#### Identifying enriched regulons in the ISCs

Transcriptional regulatory networks were constructed using Reconstruction of Transcriptional regulatory Networks and analysis of regulons (RTN) algorithm (26). 1600 human master transcription factors (27) associated target network was constructed using R package RTN with BH-adjusted Pvalue 0.05, while unstable associations were removed by bootstrap analysis (1000 resampling, consensus bootstrap > 95%). Finally, regulon activity scores were calculated by two-tailed Gene Set Enrichment Analysis. The regulon scores enriched for each subtype were fetched using Limma.

#### Identifying enriched hallmarks and immune pathways in the ISCs

Gene set enrichment analysis (GSEA) was carried out in R using the clusterProfiler package (v4.17) (28). Hallmark and ImmuneSigDB gene sets from Molecular Signatures Database (MSigDB v7.4.1) were obtained via the msigdbr package (v10.0.2) (23,25). All DE genes per ISC were ranked by their log₂ fold-change from differential expression analysis. GSEA was performed with the GSEA() function, using 1,000 gene-set permutations, a weighted scoring scheme, and gene-set size filters of 5–500 genes. Enrichment results were adjusted for multiple testing, and gene sets with a false discovery rate (FDR) < 0.05 were considered significant.

#### Cell type enrichment analysis across ISCs

To characterize the tumor microenvironment across transcriptionally defined subgroups, we applied the xCell algorithm (29) to log2-transformed TPM expression values. xCell was run using the R package xCell (version 1.1.0) with default settings, yielding enrichment scores for 67 immune and stromal cell types (including three summary score signatures) per sample.

Enrichment scores were combined with All Samples Category clusters to evaluate differences in cell type enrichment across ISC clusters. For each cell type signature, the median xCell enrichment score was calculated within each cluster and then ranked from highest to lowest to determine a relative median rank (1 being highest, 4 the lowest).

### Survival analyses

Survival analyses of baseline naïve biopsies were performed by Cox proportional hazards modeling to evaluate treatment effects on progression free survival (PFS) within HNSCC subtypes. PFS for both first-line (1L) and post-metastasis therapies (M1L) was calculated according to RECIST criteria (30). Time-to-event variables were constructed using the treatment start date of the first line of therapy as the index date. A PFS event or initiation of the next line of therapy was considered if it occurred more than 14 days after the treatment start date and within the exposure window, defined as up to two years from the start date or 30 days after the curated last known date, whichever occurred earlier. Patients were censored at the earliest of two years after treatment initiation or the curated last known date.

#### Processing public single-cell sequencing studies

Two publicly available single-cell datasets (CNP0001341 (31) and GSE195832 (32)), containing pre– and post-treatment samples from Chemotherapy and CPI, were analyzed. To assign cell type labels, both datasets were first aligned to an internal pan-cancer reference landscape using the scvi-tools (33). Once annotated, the datasets were integrated using scvi-tools, treating the study of origin as the batch variable. Cell type annotation was based on single-cell reference data provided by Nebion AG (https://www.immunai.com/), used under a service agreement that permits publication of analyses derived from this reference.

#### Assigning subtypes to samples in single-cell studies

Top 50 genes specific to each subtype were used to assign subtypes to the samples in single cells studies. Signature scores for each cell across all subtypes were calculated using Scanpy’s *score_genes* function. To account for variation across cells, we further normalized the scores using z-score normalization *(sklearn.preprocessing.StandardScaler)*. Each cell was then assigned a z-score-based predicted subtype based on the highest normalized score. The scores were further aggregated at the sample level. For each sample, the mean z-score per subtype was calculated, and the subtype with the highest average z-score was defined as the dominant subtype for that sample.

### Statistical analyses

All statistical analyses were conducted in R (V4.4.1). A wilcoxon rank-sum test was utilized for continuous data. A Fisher exact test was used for comparisons of categorical variables.

## Results

Multi-omics clustering of HNSCC via the integration of transcriptomics and genomics data using the meta-SNF algorithm followed by spectral clustering revealed the presence of four distinct ISCs; consistent across three sample categories: ISC-N, ISC-NT, and ISC (**Figure 1A**). The three sample categories yielded a high Adjusted Rand Index (ARI) of 0.98, indicating strong concordance. The term *ISC* is used throughout to refer to the consolidated immune subtype clusters that encompass both ISC-N and ISC-NT groups, given the incremental nature of the sample categories.

**Figure 1.**
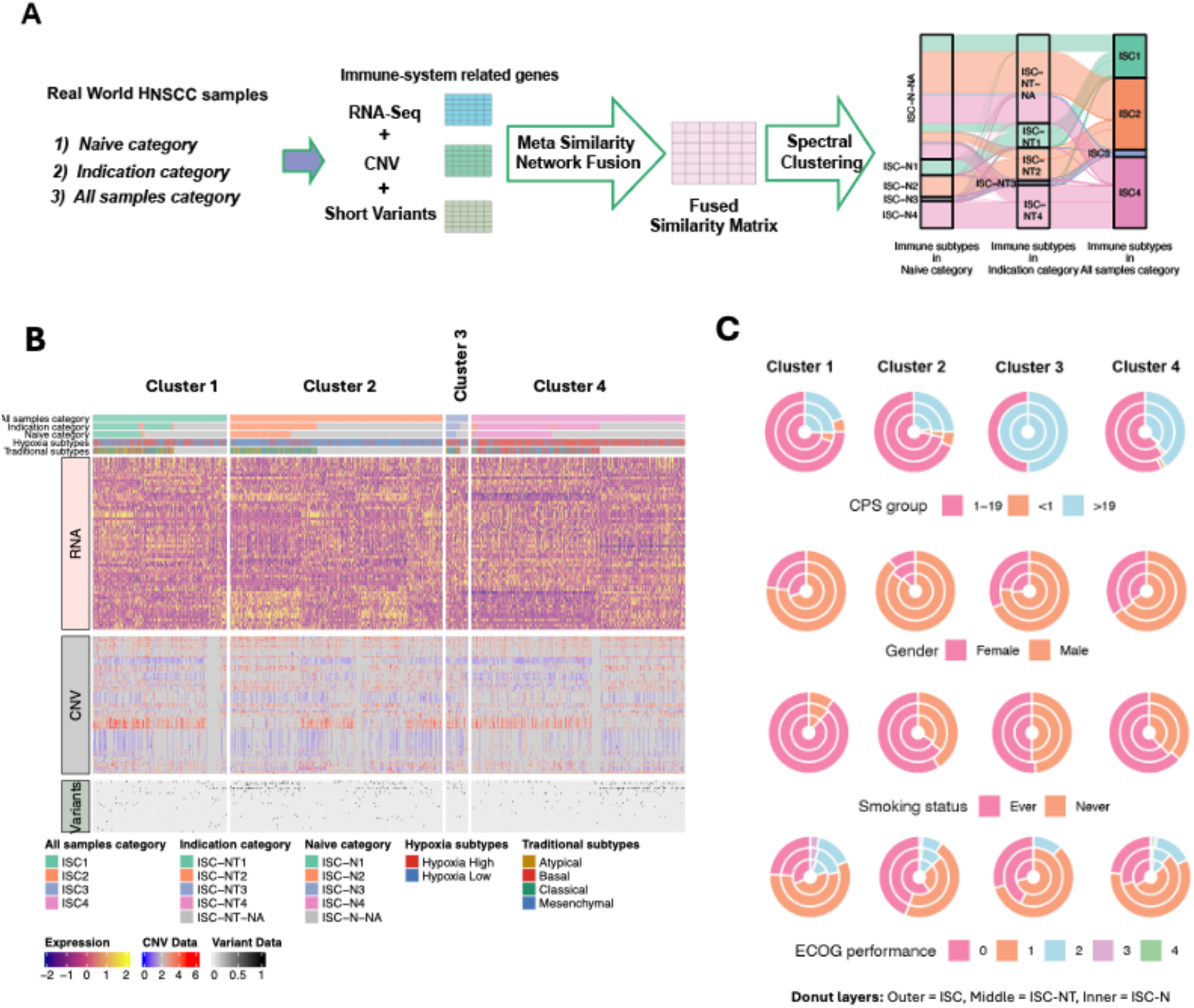
Integration of RNA, CNV, and short variant data reveals four distinct ISCs in real-world HNSCC samples. **(A)** Schematic overview of the identification of ISC using metaSNF. RNA-Seq, copy number variation (CNV), and short variant data related to immune system genes were extracted from real-world HNSCC samples, categorized into three analysis groups: Naive category, Indication category, and All samples category. A Meta Similarity Network Fusion approach was used to integrate multi-omic data into a unified similarity matrix, which was then subjected to spectral clustering to identify robust immune ISCs. The Sankey diagram on the right shows the concordance of ISC assignments across the three categories, highlighting four major clusters (ISC1–ISC4). **(B)** Overview of ISC patient groups. The heatmap shows characterization of four ISC groups based on their enrichments for different genomic and transcriptomic features. **(C)** Donut plots summarizing clinical and molecular annotations across the four ISC clusters. Each donut plot represents the distribution of a clinical variable across three annotation layers: outer = ISC (all samples), middle = ISC–NT (indication category), inner = ISC–N (naive category).

To understand the biological and clinical relevance of the ISCs, we examined their molecular, epidemiological, and clinicopathological characteristics. Patients within each cluster demonstrated unique molecular profiles, with Cluster 3 enriched for individuals with high PD-L1 CPS, suggesting potential responsiveness to CPI therapies (**Figure 1B–C, Supplementary Figure S3A–B, and Supplementary Table S1**). Age distributions varied, with Clusters 1 and 2 primarily composed of patients aged 50–70 years, while Clusters 3 and 4 included more patients aged 49 years or younger. Cluster 1 had a higher proportion of smokers and patients with elevated ECOG scores, indicative of poorer functional status. Most patients across clusters were stage 4 with “Progressive disease”. Cluster 3 contained a higher number of treatment-naïve samples in ISC-N category, while the ISCs had a balanced distribution of naïve and treated samples. Differences in visceral metastases were also observed, with lung and lymph nodes being the most common sites overall. However, Cluster 2 showed increased metastases in esophageal and digestive organs, whereas Cluster 3 had a higher prevalence of skin metastases. HPV status and sex further distinguished clusters: Cluster 2 had higher proportion of HPV-positive cases and male patients, whereas Cluster 1 was predominantly HPV-negative. HPV-positivity is associated with improved survival in HNSCC patients (34). Tumor anatomical origin varied as well, with oral cavity tumors more frequently observed in Clusters 3 and 4, and pharyngeal tumors more common in Clusters 1 and 2. Collectively, these findings support the biological distinctiveness of the four ISC groups and their potential clinical relevance in HNSCC.

To further contextualize the ISCs, we compared them against known molecular classifications, including hypoxia (defined using a curated set of 198 hallmark hypoxia genes) and traditional subtypes (i.e. basal, classical, atypical, and mesenchymal; derived from clustering the 2500 most variant genes) (**Supplementary Figure S4**). Mapping the ISC groups onto hypoxia subtypes revealed that ISC1 and ISC3 exhibited a mixed hypoxia phenotype, ISC2 was predominantly hypoxia-low, and ISC4 was characterized by a hypoxia-high profile (**Supplementary Figure S4A**). In contrast, ISC clusters did not align neatly with the traditional subtypes. ISC1 included a mixture of atypical, basal, and classical profiles; ISC2 spanned atypical and classical; ISC3 included classical and mesenchymal; and ISC4 was composed of basal and mesenchymal features (**Supplementary Figure S4B**). These observations suggest that while hypoxia patterns may partially distinguish ISC clusters, traditional subtype classifications do not fully capture the immune-based stratification revealed by ISCs, highlighting that immune subtyping captures distinct axes of tumor biology.

### Distinct genomic alteration and copy number variation among ISCs

To better understand the genomic foundations of the ISCs, we examined both mutational profiles and CNVs across the groups. Fisher’s exact test identified non-synonymous mutations significantly overrepresented in specific clusters (**Supplementary Table S2**). The overall mutational landscape was consistent with prior reports (35), with *TP53*, *PIK3CA*, *CDKN2A*, *FAT1*, and *NOTCH1* among the most frequently altered genes (**Figure 2A**). ISC2 showed a higher prevalence of *PIK3CA* and *KMT2D* mutations while *TP53* mutations were significantly enriched in ISC1 and ISC4. In ISC4, TERT promoter mutations (*TERT* 124C>T and *TERT* 146C>T) were more frequent (**Figure 2B**) and have been reported in other cancer types (36) and associated with poor prognosis in glioblastoma (37). TERT promoter mutations were further enriched in Hypoxia high subtype (**Figure 2C**). Premature stop codons in *CDKN2A* (Arg58* and Arg80*), resulted in the loss of functional p16åINK4a protein, a critical regulator of the G1-S phase transition in the cell cycle (38), were more frequently observed in ISC4 and the Hypoxia-high subtype. This enrichment may reflect converging mechanisms of tumor progression characterized by impaired cell cycle control and a hypoxic microenvironment.

**Figure 2:**
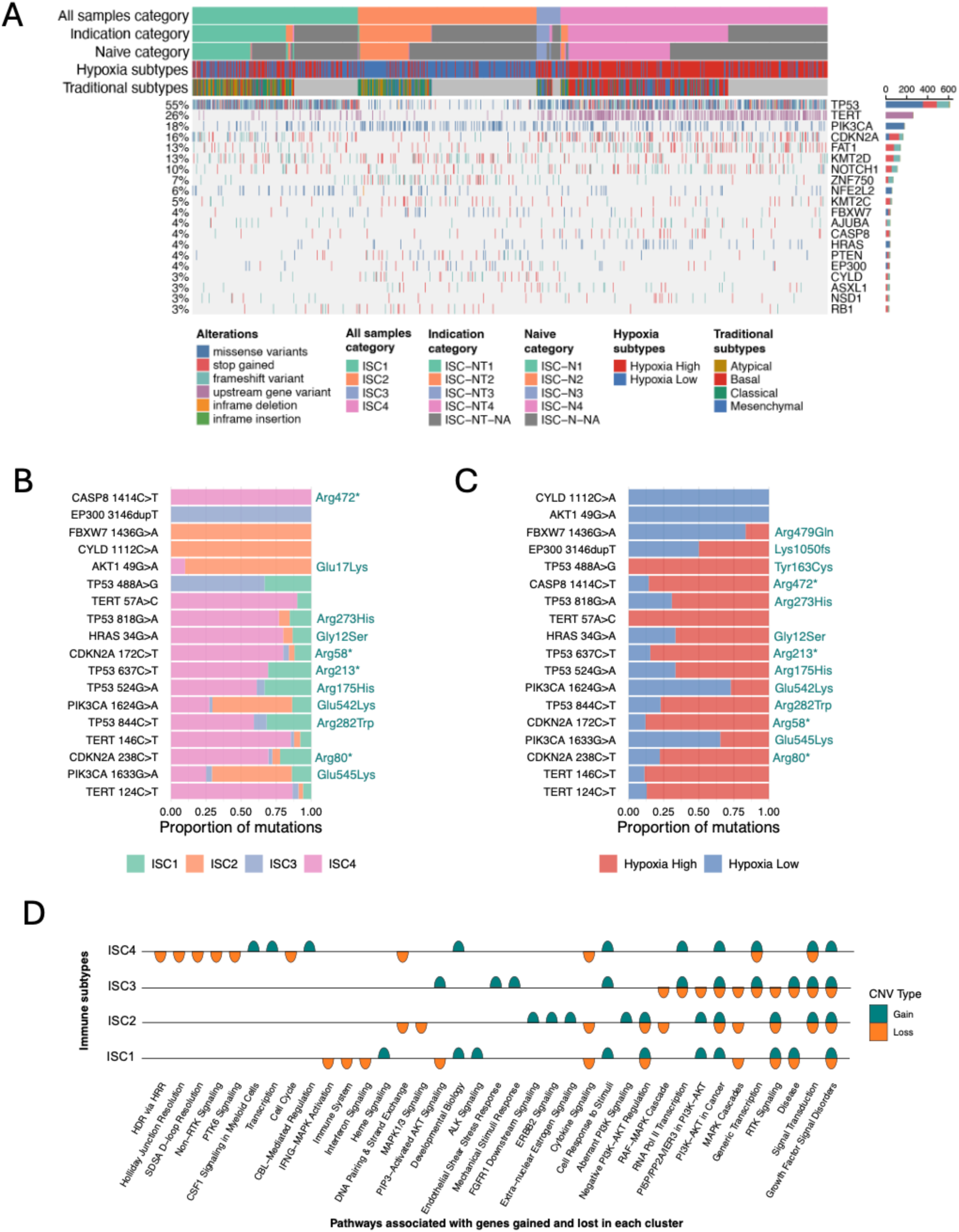
Genomic landscape of ISCs reveals subtype-specific mutations and copy number alterations. **(A)** Oncoplot showing the distribution of top 20 frequently mutated genes across immune subtype clusters (ISC1–ISC4). A horizontal barplot (right) summarizes the total frequency of each mutated gene across the cohort. **(B)** Proportion of samples within each immune subtype harboring selected non-synonymous mutations. **(C)** Proportion of samples with the same set of mutations stratified by hypoxia status (Hypoxia High vs. Low). **(D)** CNV analysis showing significant pathways (FDR < 0.05) with gene gains (green) or losses (orange) in each ISC cluster.

Complementing these findings, CNV analysis revealed distinct patterns of genomic gains and losses across the clusters (**Supplementary Figure S5, Supplementary Table S3**). All ISCs showed amplification of genes in intracellular signaling pathways, most notably within the PI3K/AKT network (**Figure 2D**), a well-established oncogenic driver (39,40). ISC1 displayed specific amplification of genes in the Signaling by ALK pathway, which was not prominent in other clusters. Conversely, deletions were enriched in pathways related to cytokine signaling, IFNG-MAPK activation, and interferon signaling highlighting potential immune evasion mechanisms. Interestingly, components of the PI3K/AKT pathway were also lost in all clusters except ISC4, indicating possible dual roles of this pathway depending on the context. Additionally, ISC2 and ISC4 showed loss of genes involved in DNA repair processes such as homologous recombination repair (HRR) and DNA pairing and strand exchange, suggesting higher genomic instability in these groups.

Together, these findings highlight distinct genomic alterations across immune subtype clusters, reflecting diverse mechanisms of tumor progression. Overall, ISC4 exhibited features of genomic instability and immune suppression, suggesting a more aggressive molecular phenotype.

### Transcriptomic analysis reveals divergent transcription and regulatory landscapes among the four immune subtypes

Using transcriptomics data, we identified differentially expressed (DE) genes between each ISC against all other samples. Among the clusters, ISC1 exhibited the fewest DE genes, whereas ISC4 had the highest number (**Supplementary Figure S6, Supplementary Table S4**). Although distinct transcriptomic and regulon activity patterns are visible in **Figure 3A**, we further integrated regulon activity (interpreted from the full transcriptome; **Supplementary Table S5**) with hallmark-level insights from Gene Set Enrichment Analysis (GSEA) of DE genes (**Figure 3B, Supplementary Table S6**) to gain a broader view of subtype-specific transcriptional programs.

**Figure 3:**
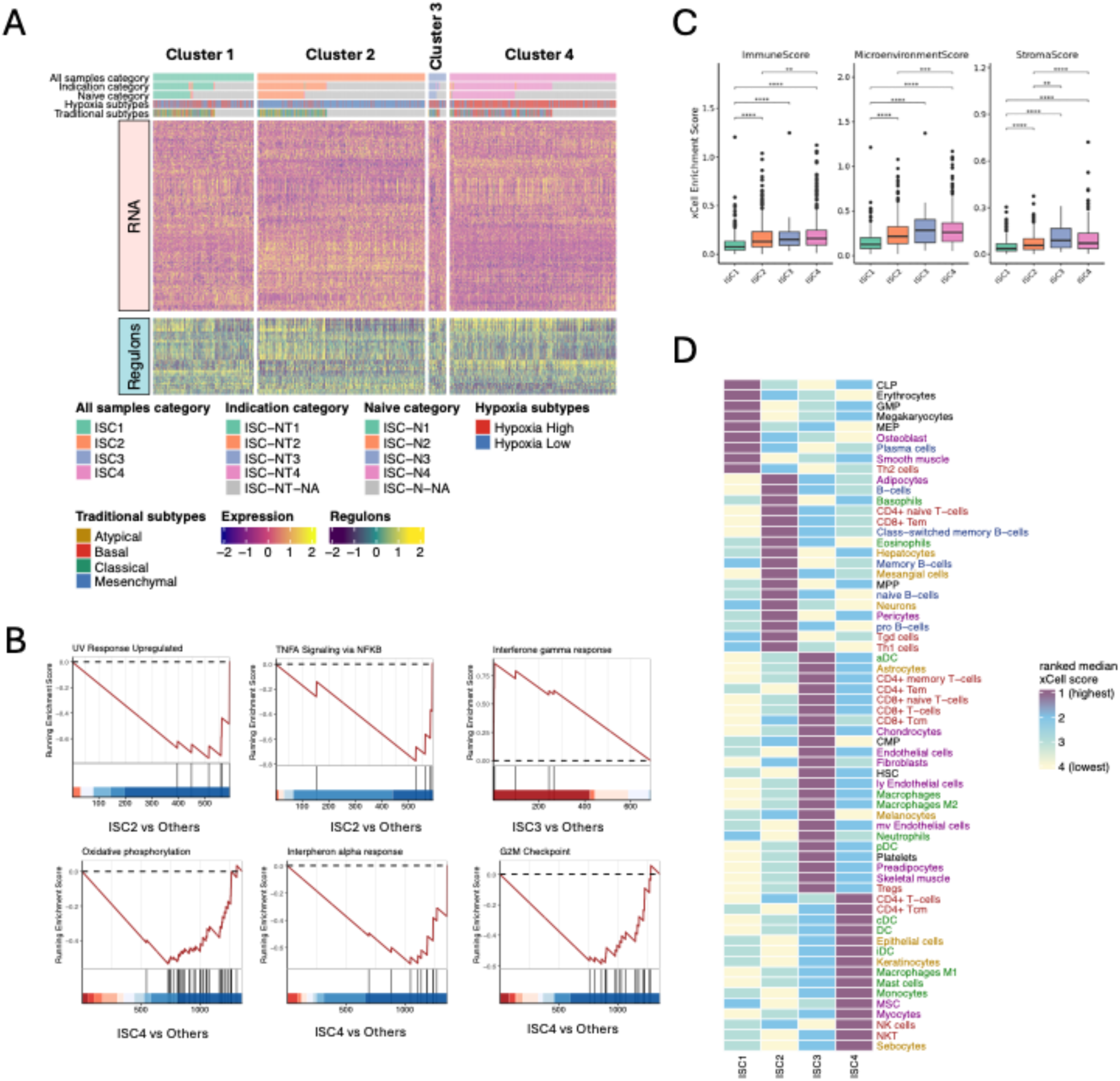
Transcriptomics and immune landscape across ISC subtypes. **(A)** Integrated clustering of RNA expression and regulon activity profiles reveals four distinct molecular clusters across all samples. **(B)** Gene set enrichment analysis highlights key hallmarks enriched in individual ISC groups compared to others. **(C)** xCell-derived summary scores i.e. ImmuneScore, MicroenvironmentScore, and StromaScore across ISC1–ISC4. Differences are statistically significant (p < 0.001, Wilcoxon test). **(D)** Heatmap of ranked median xCell enrichment scores for cell types across ISC groups.

ISC1 exhibited upregulation of Transcription factors (TFs) such as *FOXL2, TEAD2, ZEB1,* and *FOXC2*, indicating an active transcriptional program promoting epithelial-to-mesenchymal transition (EMT) and tumor plasticity (41,42). Concurrent downregulation of epithelial-associated TFs (*ELF5, GRHL3, POU2F3*) and immune regulators (*IKZF2, IRF5*) suggests loss of epithelial identity and immune exclusion, consistent with a dedifferentiated tumor phenotype (43,44). Although no pathway-level enrichments were detected by GSEA, the regulon activity highlights a regulatory landscape permissive to plasticity and immune evasion.

ISC2 showed upregulation of *CHCHD3*, pointing to activation of transcriptional and mitochondrial processes (45), yet GSEA revealed downregulation of cell cycle (E2F targets, G2M checkpoint), oxidative phosphorylation, and immune pathways (interferon gamma, TNFα/NF-κB). This suggests that while *CHCHD3* may support mitochondrial organization, the dominant phenotype is functional suppression across proliferation, metabolism, and immunity. Upregulation of EMT and KRAS signaling in GSEA complements the metabolic shift and may represent compensatory survival and invasive programs in the context of regulatory repression.

ISC3 was marked by upregulation of TFs such as *IRF5, USF1, GRHL3, PAX1*, and *BARX2*, consistent with enhanced immune regulation, developmental signaling, and epithelial identity (46,47). This aligned with GSEA findings, which showed strong interferon gamma response enrichment, supporting a more immunologically active phenotype. The presence of both EMT– and epithelial-associated TFs (e.g., *GRHL3, OVOL1*) may reflect a hybrid EMT state with preserved immune responsiveness.

ISC4 revealed enrichment of TFs including *SKIL, CSRNP3,* and *HBP1*, linked to transcriptional repression and chromatin remodeling (48). Downregulation of additional TFs involved in immune regulation (*USF1*, *PAX1*) and chromatin modifiers (*DOT1L*, *SAFB*) reinforces a broadly repressed transcriptional state (48). This was supported by GSEA, which showed widespread suppression of proliferative (E2F, G2M), metabolic (oxidative phosphorylation, myogenesis), and immune pathways (interferon responses). Reduced apical junction signaling further indicates loss of adhesion and metastatic potential.

Collectively, these findings demonstrate subtype-specific regulatory and functional programs, with concordance between transcription factor activity and downstream pathway modulation. Overlaps in TF usage across subtypes point to context-dependent roles, while the interplay between EMT, immune signaling, cell cycle control, and chromatin regulation highlights the layered complexity of transcriptional reprogramming in tumor subtypes.

### Immune Subtypes defined by their unique immune landscapes

To characterize the immune and stromal landscape across the ISCs, we integrated cell-type estimates using xCell (**Supplementary Table S7**) with enrichment analysis of immune-related gene sets from the immunesigDB (C7 collection) (**Supplementary Table S8**).

ISC1 displayed the lowest immune and stromal scores across all subtypes (**Figure 3C**). It was characterized by lower enrichment of CD8+ T cell memory, central and effector memory T cells, regulatory T cells, and B cell subsets (**Figure 3D**). Myeloid components such as dendritic cells, monocytes, and macrophages also showed lower scores as compared to other subtypes. Among stromal features, fibroblasts, endothelial cells, and skeletal muscle were low, whereas smooth muscle and common lymphoid progenitors (CLPs) were elevated.

ISC2 exhibited the highest immune score and was enriched for multiple immune cell types, including memory B cells, plasma cells, gamma-delta T cells, CD8+ effector memory T cells, and myeloid dendritic cells. Despite this, ISC2 showed relatively low levels of central and effector CD4+ T cell memory, and regulatory T cells, stromal cell types, including fibroblasts, mesenchymal stem cells, endothelial cells, and myocytes. Enrichment analysis from immunesigDB revealed enrichment of signatures associated with memory/effector CD8+ T cells and early immune response.

ISC3 showed a more balanced immune profile with elevated levels of central and effector CD4+ T cells, Tregs, and myeloid cell populations including monocytes, macrophages, and dendritic cells. It also had the highest levels of stromal and endothelial markers, along with fibroblasts and skeletal muscle. Hematopoietic stem cells (HSCs) were also relatively high in this group. GSEA enrichment analysis indicated strong MHC/HLA-mediated antigen presentation, CD8+ cytotoxic/memory T cell activity, CD4+ helper T cell enrichment indicating potential of T-cell engagement and pro-inflammatory phenotype.

ISC4 was characterized by its elevated levels of regulatory and myeloid immune cells, including Tregs, dendritic cells, neutrophils, monocytes, and macrophages. Stromal cell enrichment was also high, particularly for fibroblasts, endothelial cells, and smooth muscle. CD8+ T cell memory and CD4+ T cell memory were present at moderate levels. GSEA analysis highlighted that the subtype represents lower helper T cell signature, lacks antigen presentation, reduced CD8+ memory T cell features representing less adaptive immune coordination and immune-excluded environment, consistent with the detected genomic alterations.

Overall, these results highlight the diverse immune and stromal compositions of the ISC subtypes. The combined xCell and immunesigDB analyses reveal that ISC1 represent limited immune engagement, ISC2 exhibit stronger immune activation or stromal involvement, ISC3 show selective enrichment of immune cell types and ISC4 represent immune-excluded environment.

### Integrated immune subtype classification reveals distinct states of immune engagement and links immune activity to PFS

The integrated multi-omics analysis across clinical, mutational, transcriptional, and immune profiles detailed the ISCs in HNSCC, capturing different states of immune engagement and suppression (**Figure 4A**).

**Figure 4:**
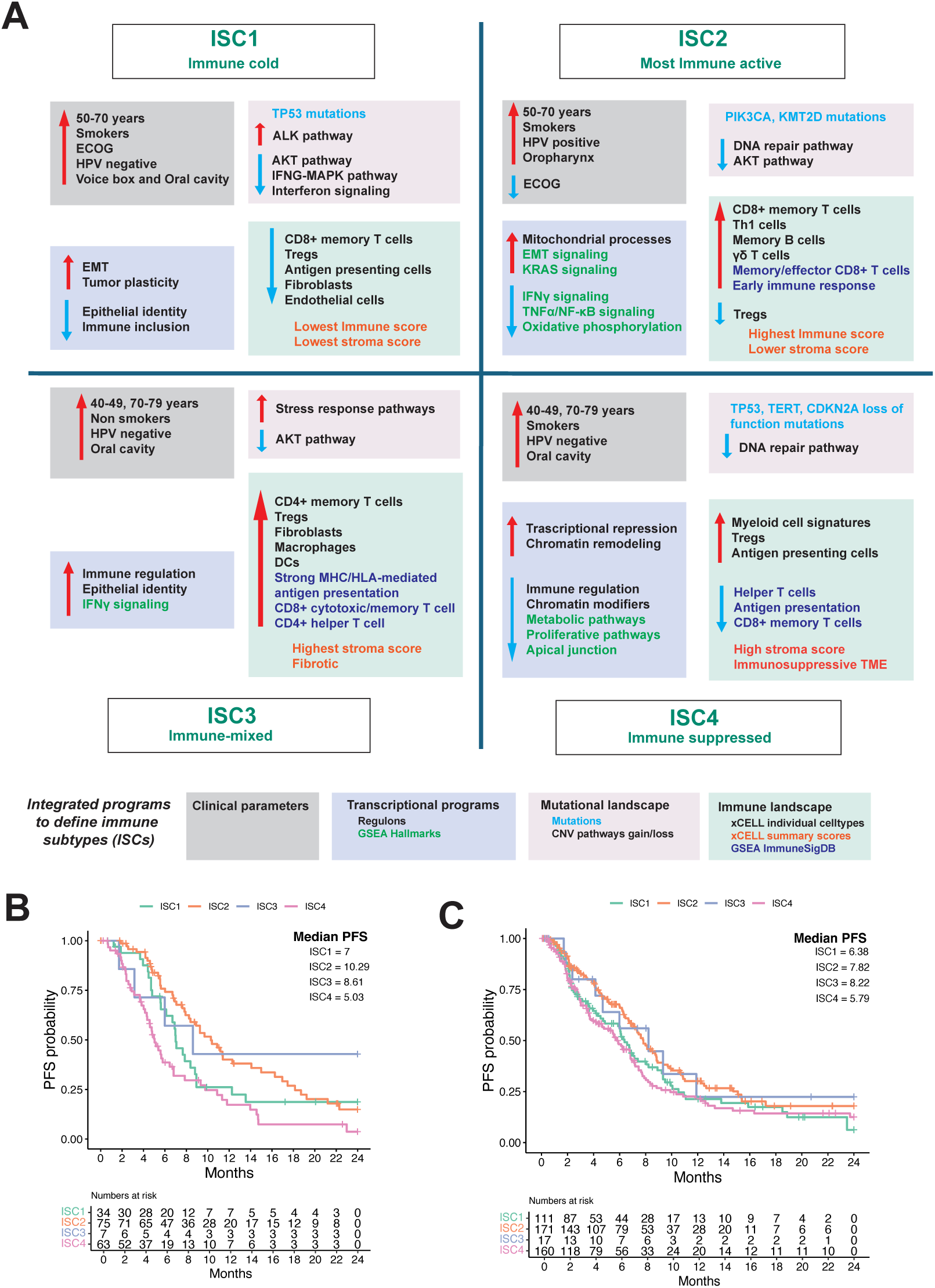
Integrated classification of ISCs (ISC1–ISC4) based on multi-dimensional features and their association with PFS. **(A)** Schematic representation of four ISCs derived from clinical, transcriptional, mutational, and immune landscape features. **(B–C)** Kaplan-Meier curves showing PFS stratified by ISC subtypes in two groups. **(B)** Progression in patients with naïve tumors with first line treatment **(C)** Progression in patients with line of treatment post metastasis or recurrence. The median PFS times are labeled for each subtype. Tables below each plot indicate the number of patients at risk over time per ISC subtype.

ISC1 represented an **immune-cold** group, marked by epithelial plasticity, immune exclusion, and low immune infiltration. ISC2 reflected the most **immune-active** state, enriched for memory and effector T cells and early immune activation pathways, although concurrent transcriptional repression signatures highlighted underlying biological complexity. ISC3 exhibited a **mixed immune** phenotype, characterized by immune regulatory programs, strong antigen presentation, and stromal remodeling, suggesting a tissue-infiltrated but functionally heterogeneous microenvironment. ISC4 was defined by **immune suppression**, genomic instability, stromal fibrosis, and enrichment of myeloid populations, consistent with a suppressed adaptive immune landscape.

While ISC2 and ISC3 both showed features of immune activation, minor discrepancies were noted between transcriptional regulon activity and xCell-derived immune cell enrichment scores, particularly in ISC3, where metabolic and EMT programs may influence immune infiltration and function. These distinctions highlight the multilayered regulation of tumor– immune interactions in HNSCC.

Kaplan-Meier analysis demonstrated that the ISC2 and ISC3 subtypes were associated with significantly improved PFS compared to ISC1 and ISC4, in both clinical settings: (1) 179 patients with naïve tumors receiving first-line therapy (1L) (**Figure 4B**) and (2) 459 patients treated after metastasis or recurrence (M1L) (**Figure 4C**). These data show that ISC classification stratifies patients by PFS across both contexts, underscoring its value as a prognostic biomarker in HNSCC.

### Subtype-specific PFS highlights immune context in treatment response

In both 1L and M1L settings, patients were stratified by the treatments they received (**Supplementary Figure S7A**). Among EGFR-mutated patients, only a small fraction received Biologic (3–7%) or Biologic + Chemotherapy (8%) treatments. Notably, Biologic + Chemotherapy was evenly distributed across all subtypes.

In the 1L setting, Chemotherapy was the most preferred treatment overall whereas in the M1L setting, it was predominantly administered to ISC4 patients (42%). Post-metastasis, CPI was more frequently used in ISC2 patients (40%). The combination of Chemo + CPI was observed in 41% of ISC2 patients and 30% of ISC4 patients.

No significant differences in median PFS were observed across treatment groups in either the 1L (**Supplementary Figure S7B**) or M1L (**Supplementary Figure S7C**) settings. To gain further insight, we stratified treatments by subtype. In the 1L setting (**Supplementary Figure S7D**), Chemo was the predominant treatment modality. Accordingly, alternative therapies are not discussed further within this context to maintain focus on the predominant clinical practice.

The M1L setting is particularly relevant in clinical practice, as many patients are diagnosed at the stage of metastasis or recurrence. While traditional molecular subtypes showed prognostic value in the 1L setting (**Supplementary Figure S8A**), they had limited utility in the M/R setting, with no significant differences in PFS (**Supplementary Figure S8B**), making them less informative for guiding treatment. In contrast, hypoxia subtypes offered both prognostic and treatment-related insights: patients with low hypoxia consistently showed better survival across both settings (**Supplementary Figure S8C and S8D**). However, they did not display subtype-specific preferences for treatment—low hypoxia was associated with better outcomes regardless of the therapy administered (**Figure 5A**).

**Figure 5:**
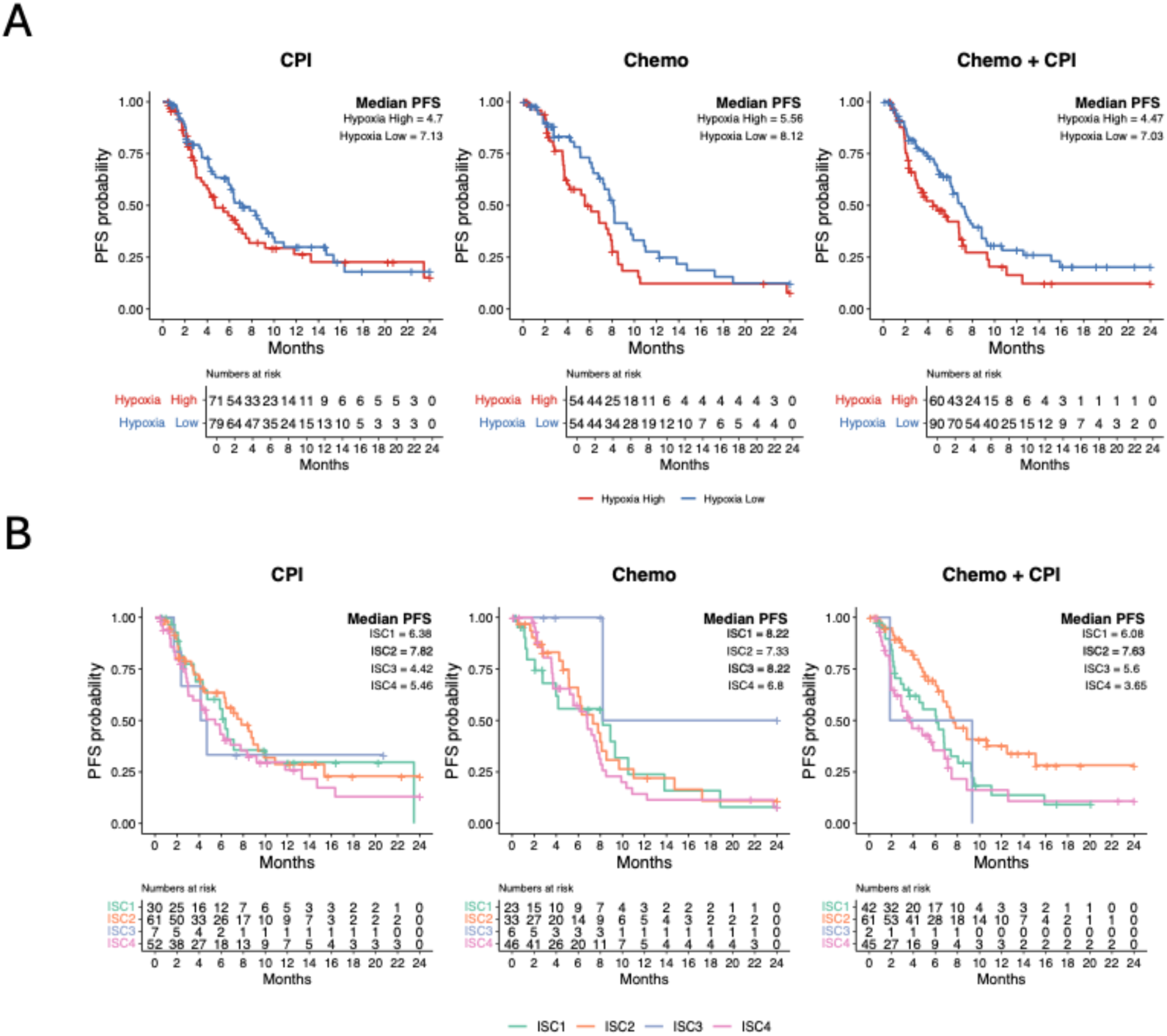
Treatment-specific PFS patterns in M1L settings in hypoxia and immune subtypes (ISCs). PFS curves for patients in the M1L setting receiving Chemo, CPI, or Chemo + CPI grouped by (A) hypoxia subtypes and (B) immune subtypes (ISCs). Median PFS and number at risk are indicated.

In contrast, the ISC classification provided even more granular clinical resolution (**Figure 5B**). The immune-active ISC2 subtype showed the strongest response to CPI and Chemo + CPI. Overall, ISC stratification identified 12% more patients in ISC2 to achieve extended survival compared to treatment-based stratification alone. ISC1 and ISC3 responded best to chemotherapy, with ISC3’s benefit likely driven by its mixed immune profile and stromal remodeling. ISC4, derived modest benefit from chemotherapy but had the poorest overall survival. These divergent outcomes underscore how ISC2 and ISC4 represent opposite ends of the immune-response spectrum i.e. active vs aggressive, respectively and illustrate the added clinical value of ISC classification over traditional or hypoxia-based models in the metastatic setting. Additionally, the added clinical value of immune-driven stratification in the M1L setting, offering refined guidance beyond traditional or hypoxia-based classifications. Notably, Biologic and Biologic + Chemotherapy groups are not shown, as these therapies were administered to pre-stratified patients and yielded the highest median PFS.

### Immune landscape dynamics reflect treatment and disease evolution in RW cohorts

To explore the evolution of ISCs across tumor progression stages, we analyzed patients with paired biopsies. Although the number of patients with longitudinal (pre– and post-progression) samples in RW cohort was limited, these cases provide valuable insight into how the tumor immune microenvironment shifts over time. To complement these findings, we also assessed paired biopsies within the single-cell HNSCC landscape (**Supplementary Figure S9**). By comparing the immune states between initial and subsequent biopsies, we could assess dynamic transitions in immune subtype alongside cell-type–level remodeling, potentially driven by prior treatments, metastatic spread, or other disease-related factors.

Longitudinal analysis of second biopsies from 44 patients revealed immune phenotype shifts in 15 cases (**Supplementary Figure S10A**). Several transitions were observed across subtypes, with immune-cold biopsies in ISC1 showing notable plasticity. Five out of 11 ISC1 biopsies shifted to other states (**Supplementary Figure S10B)**, of which four transitioned to immune suppressed state i.e. ISC4, indicating a predominant trajectory toward immune suppression. Mixed phenotype samples in ISC3 showed heterogeneous outcomes, transitioning to both immune active (ISC2) and suppressed (ISC4) states, suggesting ISC3 represents an intermediate immune profile. In an independent cohort with single-cell data available for both pre– and post-treatment samples following chemotherapy and CPI, we observed a consistent trend. Most of these samples were classified as ISC1 and ISC3 subtypes, with relatively few ISC2 and ISC4 cases (**Supplementary Figure S10C**).

Among patients who exhibited a change in immune subtype following treatment, we observed corresponding shifts in the composition of the tumor microenvironment revealing distinct cellular compartment patterns between biopsies (**Figure 6A**). For example, transitions to the immune suppressed (ISC4) phenotype showed greater variability in ImmuneScore, whereas transitions to immune-active (ISC2) state were associated with shifts in stromal components. The individual summary scores revealed distinct trends depending on the direction of immune subtype transition. Transitions from ISC1 to ISC4 were frequently associated with increases in the summary scores, whereas transitions from ISC3 to ISC4 generally showed decreases (**Figure 6B**). These patterns indicate that the impact of subtype switching on the tumor microenvironment is direction dependent. Additionally, the patient-level trajectories highlighted substantial heterogeneity within each transition group, suggesting that even among patients undergoing the same subtype shift, the extent and nature of cellular remodeling can vary considerably.

**Figure 6.**
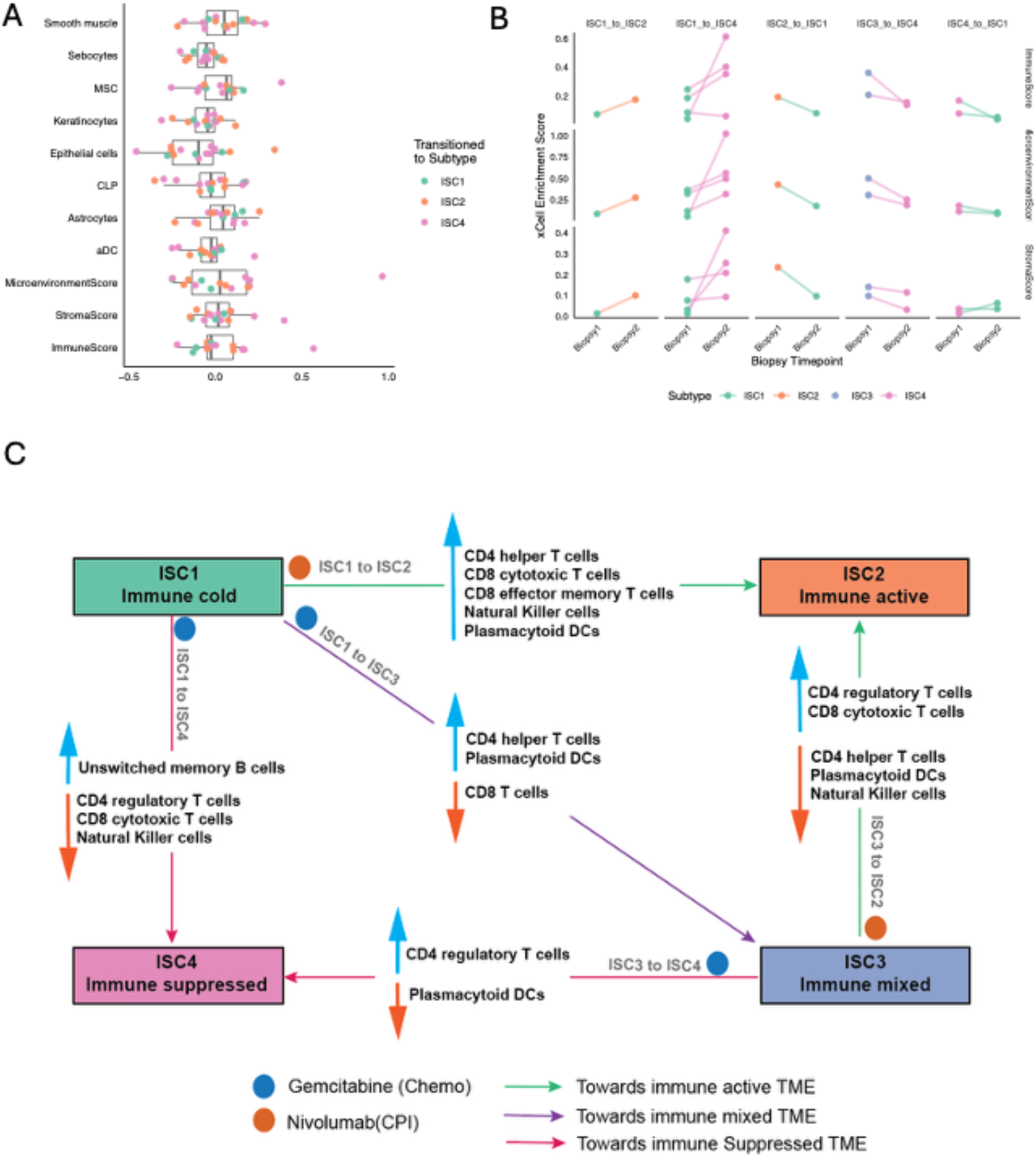
Immune subtype transitions and corresponding tumor microenvironment remodeling. **(A)** Changes in xCell enrichment scores for top tumor microenvironmental cell types between paired biopsies, stratified by final immune subtype. The top 10 signatures shown were selected based on the largest median change in score across all patients. **(B)** Directional changes in ImmuneScore, MicroenvironmentScore, and StromaScore by specific immune subtype transition. Each line connects a patient’s pre– and post-biopsy scores, colored by resulting immune subtype. **(C)** Schematic representation of immune subtype transitions in response to treatment. The transitions are annotated by changes in immune cell proportions. Upregulated (blue) and downregulated (orange) immune cell types are derived from paired single-cell analyses (see Supplementary Figure S11).

To understand the cellular basis of these bulk-level trends, we leveraged the single-cell resolution of the dataset to annotate each transition with the direction of change in key immune populations, revealing reproducible patterns of immune remodeling across subtype shifts (**Figure 6C**; **Supplementary Figure S11**). Transitions from ISC1 (Immune Cold) to more inflamed states such as ISC2 (Immune Active) or ISC3 (Immune Mixed) were associated with increased infiltration of CD4 helper T cells, CD8 cytotoxic and effector memory T cells, natural killer (NK) cells, and plasmacytoid dendritic cells (pDCs), reflecting a shift from immune exclusion to immune engagement.

In contrast, transitions into ISC4 (Immune Suppressed), especially from ISC1 or ISC3, were marked by increases in regulatory T cells (Tregs) and unswitched memory B cells, alongside reductions in CD8 T cells, NK cells, and pDCs, consistent with a more tolerogenic immune landscape. Transitions from ISC2 to ISC3 were characterized by selective immune dampening, with loss of CD4 helper T cells, pDCs, and NK cells despite retention of CD8 T cells and Tregs, indicating functional modulation rather than complete immune loss. Finally, ISC3→ISC4 transitions showed further suppression, with consistent downregulation of pDCs and persistence of Tregs, highlighting a trajectory toward immune exclusion and stromal dominance.

Notably, these subtype transitions showed strong treatment-specific patterns. Transitions toward the immune-active state were observed exclusively in patients treated with checkpoint inhibitors, suggesting CPI drives immune engagement. In contrast, transitions to immune suppression state were seen only in chemotherapy-treated patients. These findings imply that the direction of immune subtype remodeling is shaped not only by intrinsic tumor evolution but also by the type of therapy administered.

Collectively, these analyses underscore that immune subtype transitions are not merely categorical shifts but reflect direction-specific, patient-variable remodeling of the tumor microenvironment anchored at two divergent poles. On one end, transitions into ISC2 (e.g., ISC1→ISC2 and ISC3→ISC2) occurred exclusively after CPI therapy, marking a CPI-driven re-engagement of effector/memory T cells, NK cells, and pDCs and delineating a favorable immune-activated trajectory. At the opposite pole, chemotherapy-associated shifts into ISC4 (notably ISC1→ISC4 and ISC3→ISC4) converged on an immune-suppressed, stromal-fibrotic landscape enriched for Tregs and memory B cells, a signature of treatment resistance and poorer outcomes. By framing ISC2 and ISC4 as the bipolar extremes of immune evolution, these findings highlight how serial monitoring of subtype conversion could guide adaptive treatment decisions and prognostication.

## Discussion

In this study, we identified four distinct ISCs in HNSCC using an integrative multi-omics framework. Leveraging transcriptomic, mutational, and copy number data from 1,102 real-world patient samples, we derived a clinically relevant classification system that captures diverse states of immune engagement and suppression. These ISCs exhibited distinct biological, genomic, and immune features and were predictive of PFS across both 1L and M1L treatment settings. Importantly, our approach preserved the full immune gene landscape and circumvented feature reduction, allowing for a comprehensive view of immune heterogeneity in HNSCC.

Our findings build on prior classification efforts such as those from TCGA datasets, which emphasized transcriptomic or molecular subtypes, and immune phenotyping frameworks that defined tumors as immune-inflamed, –excluded, or –desert (11). Unlike previous studies, which were typically limited to controlled cohorts with relatively homogeneous populations, our analysis included a broad and diverse RW dataset with varied prior treatment histories. This diversity revealed subtypes not fully captured by traditional classification systems and highlighted the interplay between immune status and hypoxia. Notably, while hypoxia subtypes partially overlapped with our ISCs, traditional molecular subtypes showed poor concordance with immune clusters. This disconnect reinforces the clinical relevance of immune-driven stratification: whereas traditional subtypes reflect intrinsic tumor biology, immune-based classification more directly informs prognosis and predicts responsiveness to immunotherapies. In a clinical setting increasingly reliant on immune-modulating treatments, immune contexture offers more actionable insights than classical stratifications.

Based on biological and clinically relevant features the subtypes could be identified as ISC1: immune-cold, EMT-enriched, ISC2: immune-active with favorable survival, *PIK3CA* and *KMT2D* mutations, ISC3: hybrid with strong antigen presentation and stromal remodeling, and ISC4: immune-suppressed, hypoxic, genomically unstable, TERT promoter region mutations, Premature stop codons in *CDKN2A*. These subtypes showed limited alignment with traditional molecular classifications but closely mapped to patterns observed in immune-based frameworks. Notably, our findings are consistent with prior reports (49), which also described distinct immune-enriched subtypes with differential inflammatory signaling and clinical outcomes. The convergence of our ISCs with independently derived immune phenotypes underscores the reproducibility and translational relevance of immune-driven stratification across tumor types and datasets.

Beyond baseline classification, we show that ISCs can shift over time in response to treatment or disease progression. Although our analysis was limited by the relatively small number of patients with longitudinal samples, we observed that these dynamic transitions align with prior studies demonstrating treatment-induced immune remodeling, particularly the immunostimulatory effects of checkpoint inhibitors (50,51) and the immunosuppressive potential of chemotherapy (52,53). Future studies incorporating spatial, longitudinal, and treatment-matched profiling will be crucial to fully capture the complexity and kinetics of immune subtype evolution.

Looking ahead, the ISC framework presents prospects to inform biomarker development in HNSCC. Identifying immune active samples could help identify candidates more likely to benefit from immune checkpoint inhibitors, offering a complement to existing biomarkers such as PD-L1 CPS (54). Conversely, immune suppressed candidates may benefit from rationally designed combination therapies targeting stromal barriers, TGF-β signaling, or hypoxia-induced immunosuppression (55,56). The observed dual role of oncogenic pathways like PI3K/AKT and TERT mutation levels across subtypes also points to context-specific vulnerabilities that could be exploited therapeutically (37,57).

Importantly, ISC2 emerges as the prototypical CPI-responsive subtype, while ISC4 defines a chemo-tolerant, immune-excluded niche with poorest outcome. The immune engagement of two different subtypes could serve as predictive biomarkers for therapy decisions. Additionally longitudinal monitoring of subtype conversion, particularly early shifts toward ISC4, could flag emerging resistance and inform timely escalation to combination or alternative regimens.

The diverse and clinically representative nature of our cohort underscores the real-world relevance of this classification system, moving beyond the limitations of trial-restricted datasets. Ultimately, integrating ISC classification into clinical workflows—potentially through automated pipelines—may enable more precise stratification for trials and treatment (58), especially in a biologically and clinically complex cancer such as HNSCC.

## Conclusions

In this large-scale, real-world analysis of HNSCC, we established a multi-omics-based immune subtyping framework that revealed four clinically and biologically distinct ISCs. These ISCs not only captured key aspects of immune activation and suppression but also showed clear associations with PFS and treatment response across diverse therapeutic contexts. By preserving the full immune gene landscape and incorporating genomic and transcriptomic features, our approach offers a more comprehensive and clinically applicable classification than existing methods. ISC2 emerged as the prototypical CPI-responsive subtype, while ISC4 defined a chemo-tolerant, immune-excluded niche associated with the poorest outcomes. The differential immune engagement across subtypes functioned as predictive biomarkers for therapy response. Longitudinal monitoring further revealed subtype conversion over time, with early shifts toward ISC4 corresponding to emerging resistance. This work underscores the potential of integrative immune subtyping to guide treatment decisions, optimize immunotherapy use, and refine future clinical trial designs. As immunotherapy continues to expand in HNSCC, ISC-based stratification may help identify patients most likely to benefit and those in need of alternative or combinatorial strategies.

## Data availability

De-identified, individual-level data used in this research was collected in a real-world healthcare setting by Tempus Inc. and is subject to controlled access for privacy and contractual reasons. The ethics committee and/or informed consent does not allow for public availability. Derived data supporting the conclusions of this article are included within the article and its additional files.

## Authors’ Contributions

**IK:** Conceptualization, data curation, formal bulk and single-cell data analysis, supervision, writing–original draft, project administration, writing–review and editing. **TdS:** Formal bulk-data analysis, writing–original draft, writing–review and editing. **SDvA:** Formal bulk-data analysis, writing–original draft, writing–review and editing. **MB:** Data curation, writing– review and editing. **FVB:** Single-cell landscape creation. **MJ:** Data curation, formal bulk-data analysis, writing–review and editing. **SS:** writing–review and editing. **BWH:** writing–review and editing. **ICRMK:** Conceptualization, supervision, writing–review and editing.

## Acknowledgments

We acknowledge the use of ChatGPT-4o for initial editing and language refinement of the manuscript. The authors subsequently performed further editing and revisions to finalize the text.

## Supplementary Figures

**Supplementary Figure S1:**
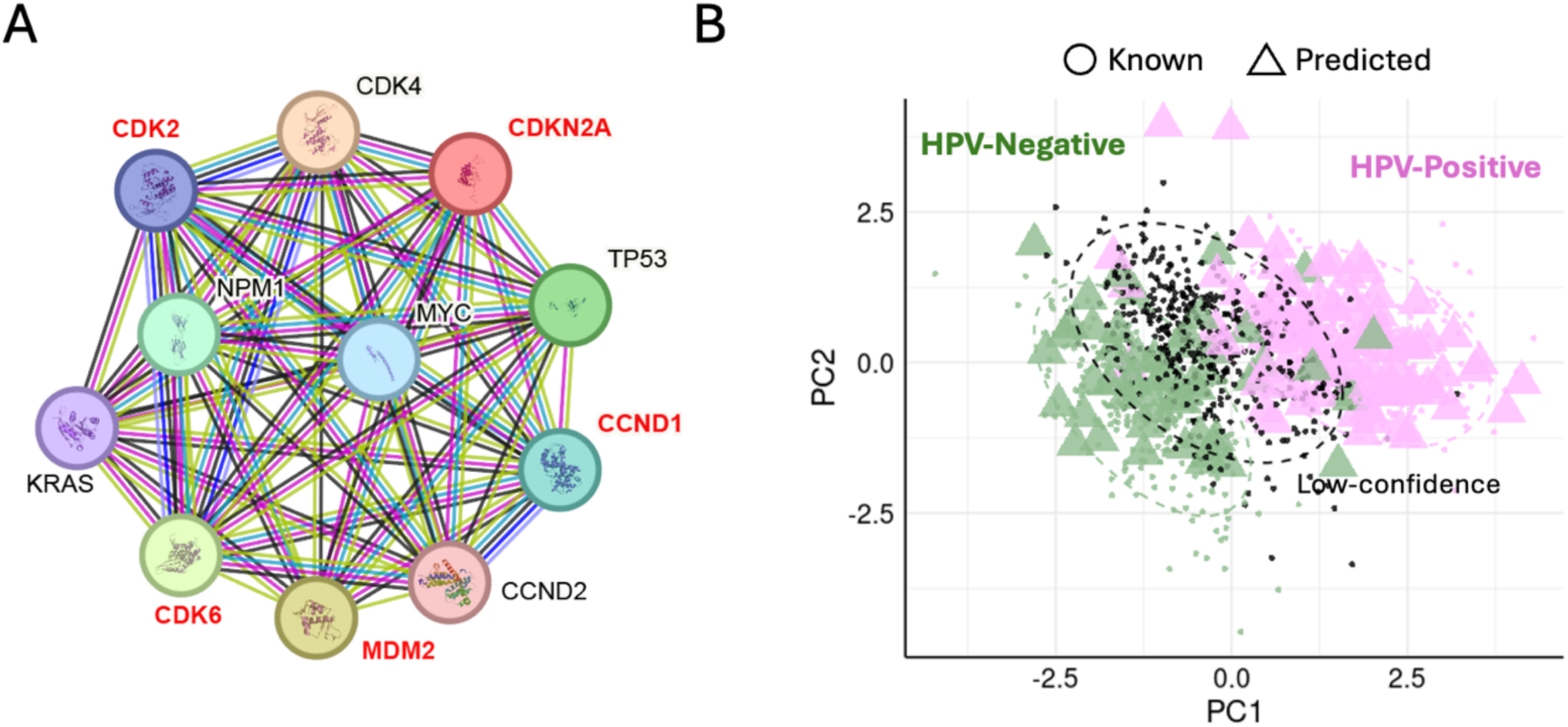
Co-expression network of genes associated with p16 (CDKN2A) and HPV status prediction for unknown samples. **(A)** Protein–protein interaction network of genes co-expressed with *CDKN2A* (p16), generated using the STRING database. Nodes represent proteins, and edges represent evidence-based associations including co-expression, experimental data, and curated databases. Genes labeled in red (e.g., *CDK2*, *CDK6*, *CCND1*, *MDM2*) were ranked higher in feature importance using a Random Forest model. **(B)** Principal component analysis (PCA) of transcriptomic data used to visualize HPV status predictions from an SVM model, based on the five red-labeled genes in panel A. Known HPV-positive (purple) and HPV-negative (green) samples were used to train the model. Triangles indicate predicted HPV status for previously unclassified samples. Samples within the dashed circle were considered low-confidence predictions and were not assigned an HPV status.

**Supplementary Figure S2:**
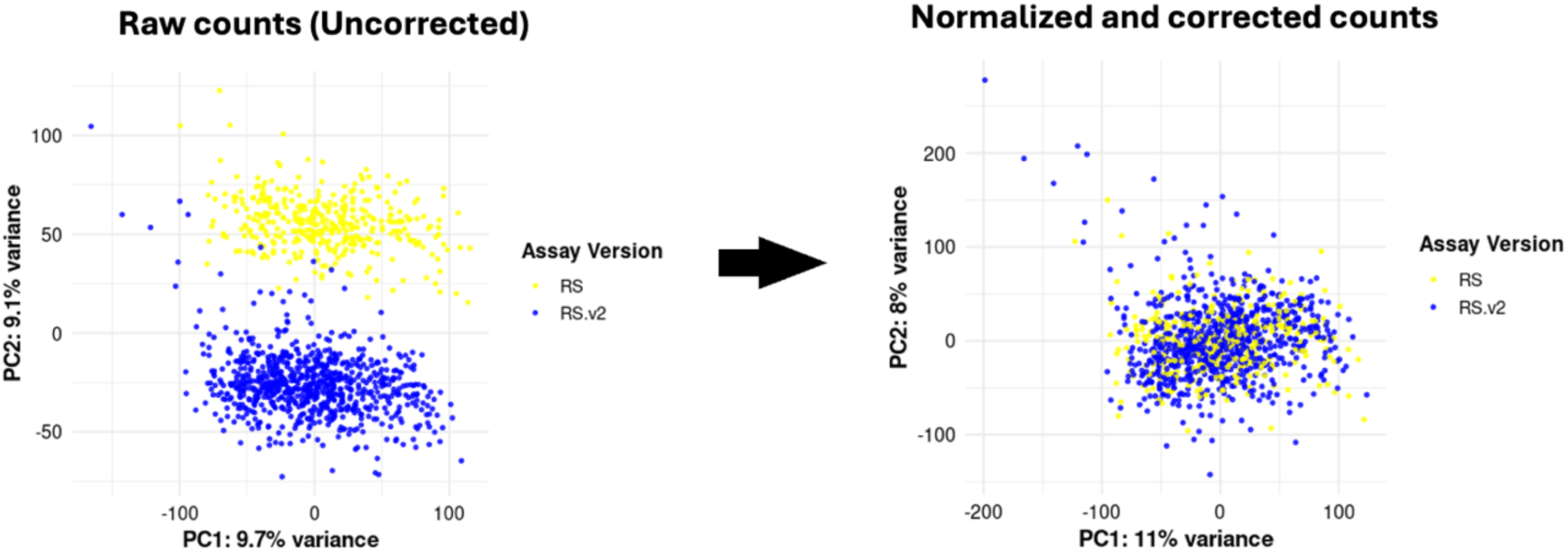
Batch correction across assay versions. Principal component analysis (PCA) of gene expression data before and after normalization and batch correction. **(Left)** Raw (uncorrected) counts show clear separation by assay version: RS (yellow) and RS.v2 (blue). **(Right)** After normalization and batch correction, samples from both assay versions are mixed, indicating successful mitigation of batch effects.

**Supplementary Figure S3:**
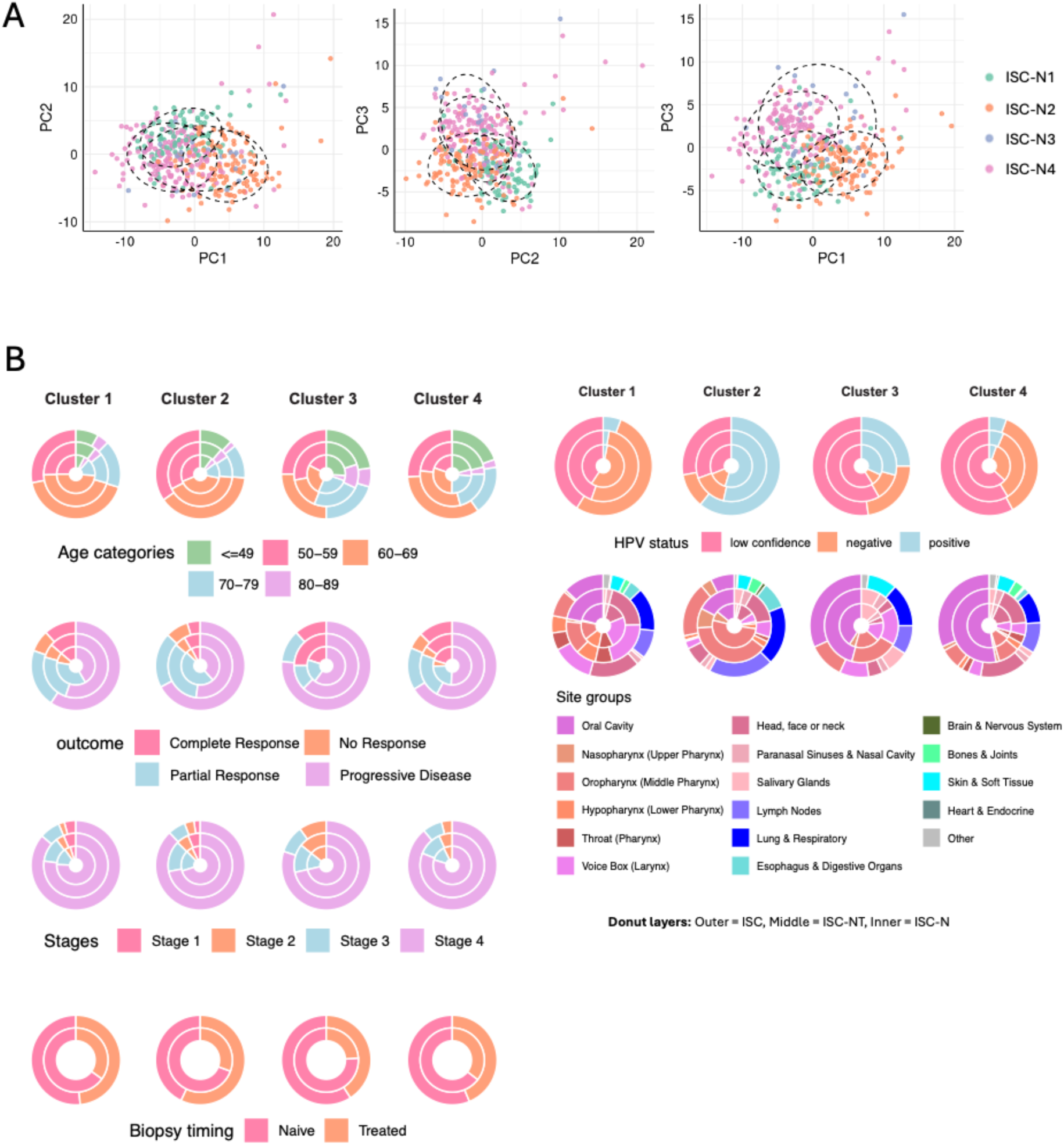
Patient stratification and clinical characteristics across immune subtype clusters. **(A)** PCA plots showing separation of immune subtype clusters (ISC-N1 to ISC-N4) in three PC combinations (PC1 vs PC2, PC2 vs PC3, and PC1 vs PC3). The dashed ellipses mark the dense core of each cluster. **(B)** Donut plots summarizing clinical and demographic characteristics across clusters. Each cluster (immune subtype group) is represented by a column of donut charts. The layers represent different data levels: outer (ISC), middle (ISC-NT), and inner (ISC-N).

**Supplementary Figure S4:**
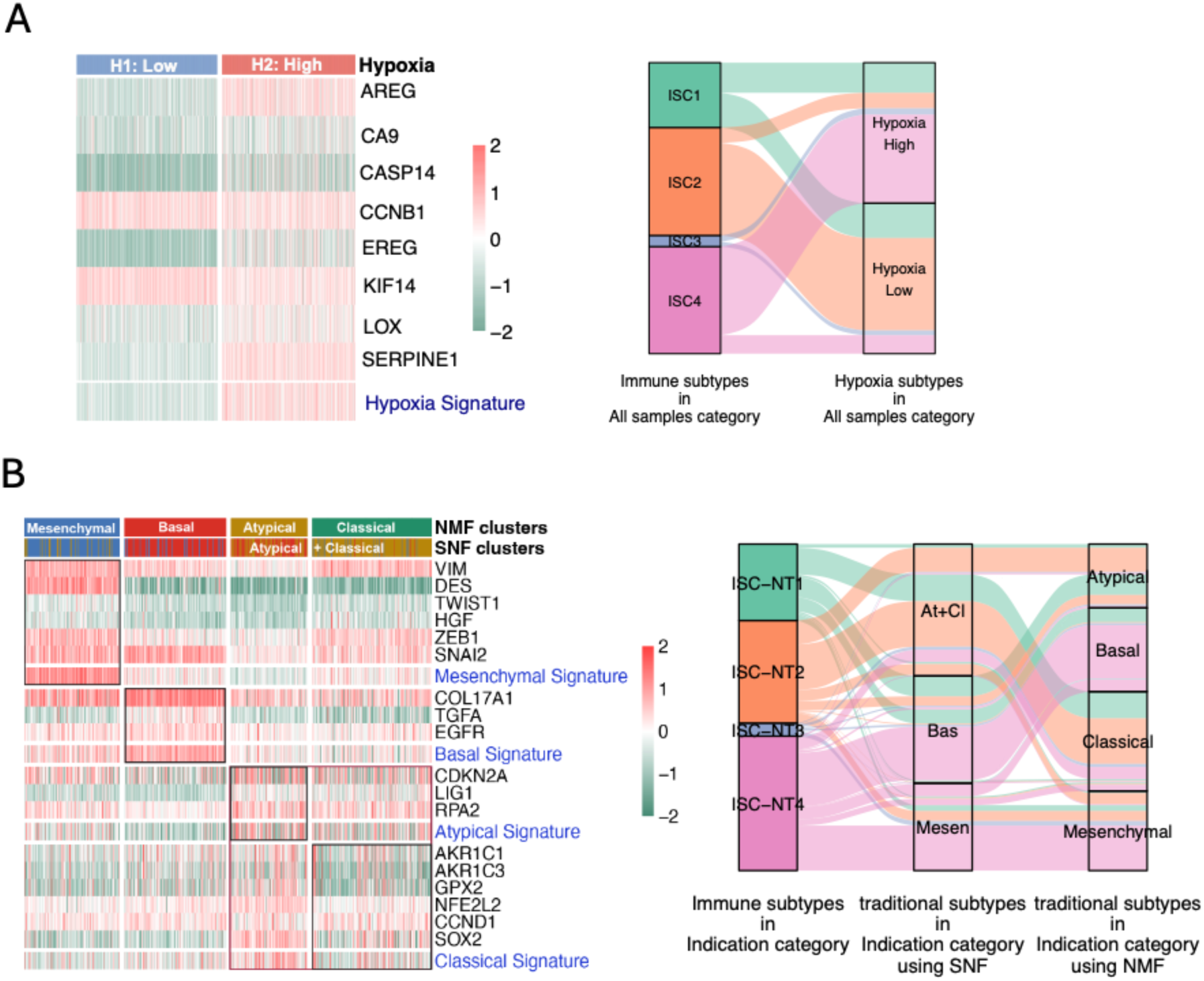
Relationship of immune subtypes with hypoxia states and traditional HNSCC subtypes. **(A)** Hypoxia subtyping: Heatmap (left) shows expression of hypoxia signature genes across samples stratified into low (H1) and high (H2) hypoxia groups. Sankey plot (right) illustrates the distribution of immune subtypes (ISC1–ISC4) across hypoxia subtypes. **(B)** Traditional HNSCC subtypes: Heatmap (left) shows expression of gene signatures corresponding to four known HNSCC subtypes (Mesenchymal, Basal, Atypical, and Classical) as defined by NMF and SNF clustering. Sankey plot (right) shows the mapping of immune subtypes (ISC-NT1 to ISC-NT4) to traditional subtypes derived using SNF and NMF methods.

**Supplementary Figure S5:**
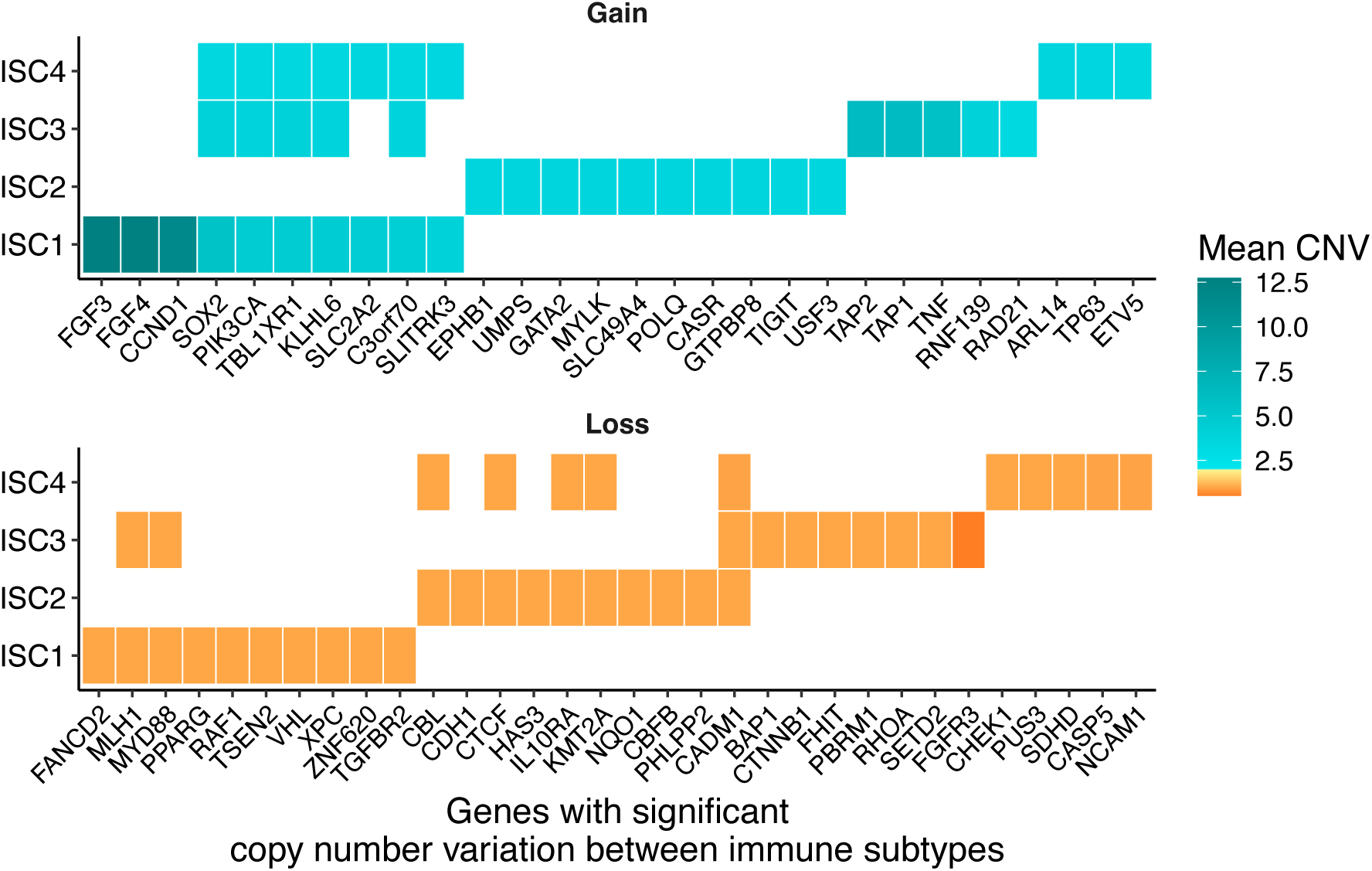
Copy number alterations in genes with significant variation across immune subtypes (ISC1–ISC4). The tile plot displays mean copy number values for genes with statistically significant CNV differences between subtypes. The top panel shows copy number gains (mean CNV > 2), and the bottom panel shows losses (mean CNV < 2). Tile color indicates the mean CNV value, transitioning from orange (loss) to teal (gain).

**Supplementary Figure S6:**
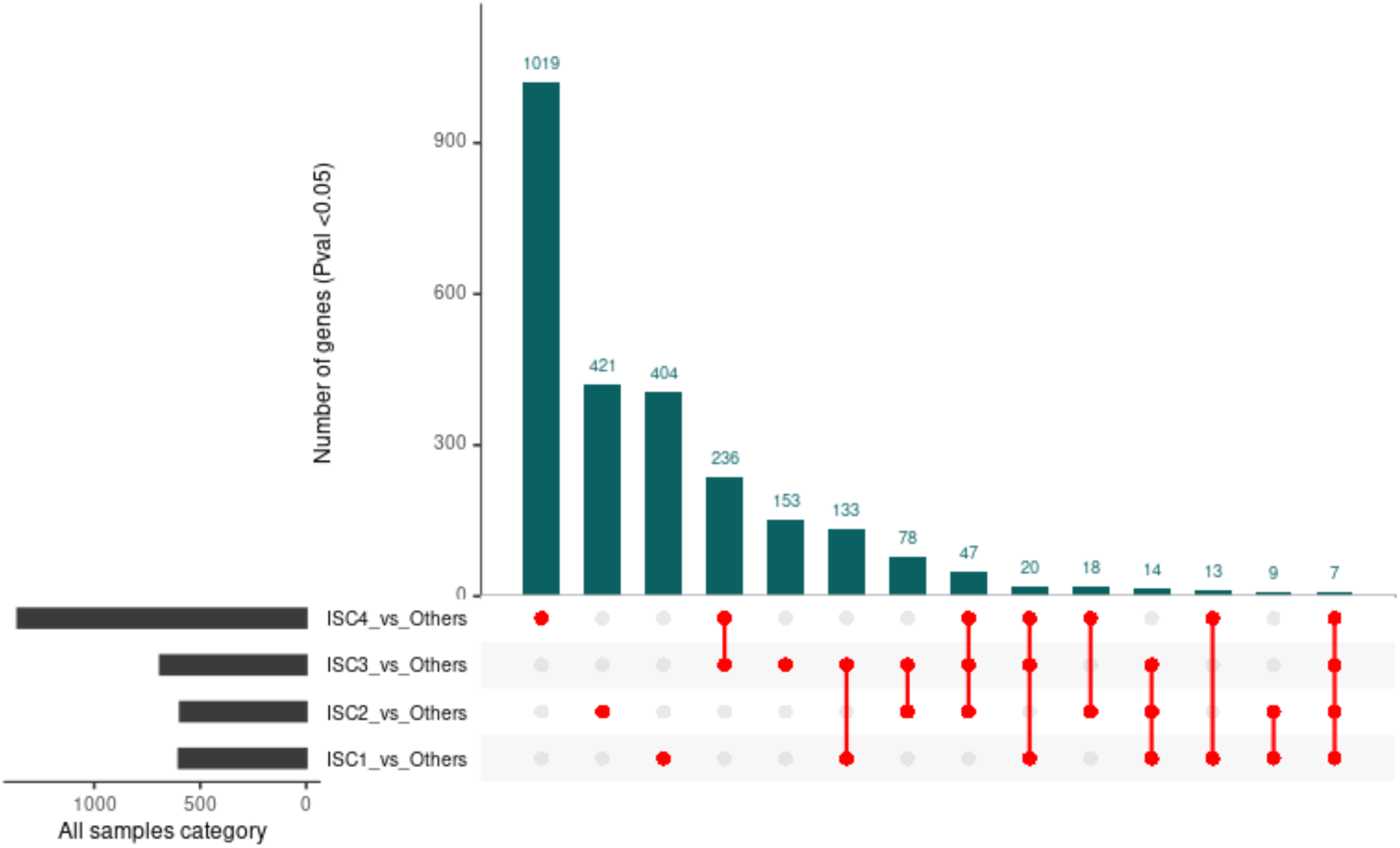
Differentially expressed (DE) genes across immune subtype clusters. Bar plot and UpSet plot showing the number of differentially expressed genes (Pvalue < 0.05) for each immune subtype cluster (ISC) compared to the rest of the cohort. The UpSet plot below illustrates the intersections of DE genes among the comparisons, indicating the degree of shared and unique transcriptional changes across the ISC clusters.

**Supplementary Figure S7:**
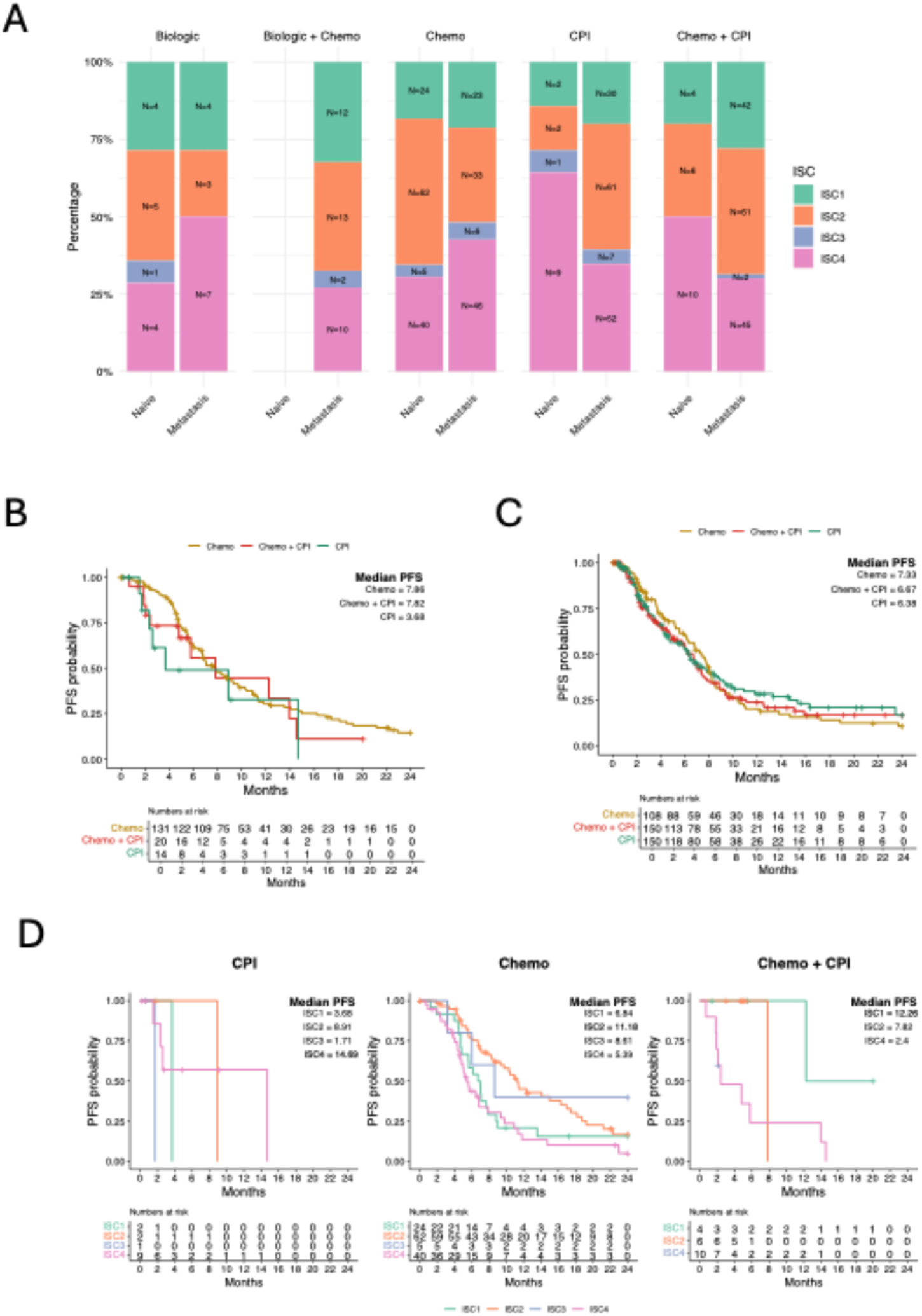
Treatment-specific progression-free survival (PFS) patterns in each subtype across clinical settings. **(A)** Distribution of treatments across ISC subtypes in both 1L (naive) and M1L (metastatic/recurrent) settings. **(B)** PFS curves for patients in the 1L setting receiving Chemo, CPI, or Chemo + CPI. **(C)** PFS curves for patients in the M1L setting receiving Chemo, CPI, or Chemo + CPI. **(D)** Treatment-specific PFS curves stratified by ISC subtype in the 1L setting. Median PFS and number at risk are indicated for all KM plots.

**Supplementary Figure S8:**
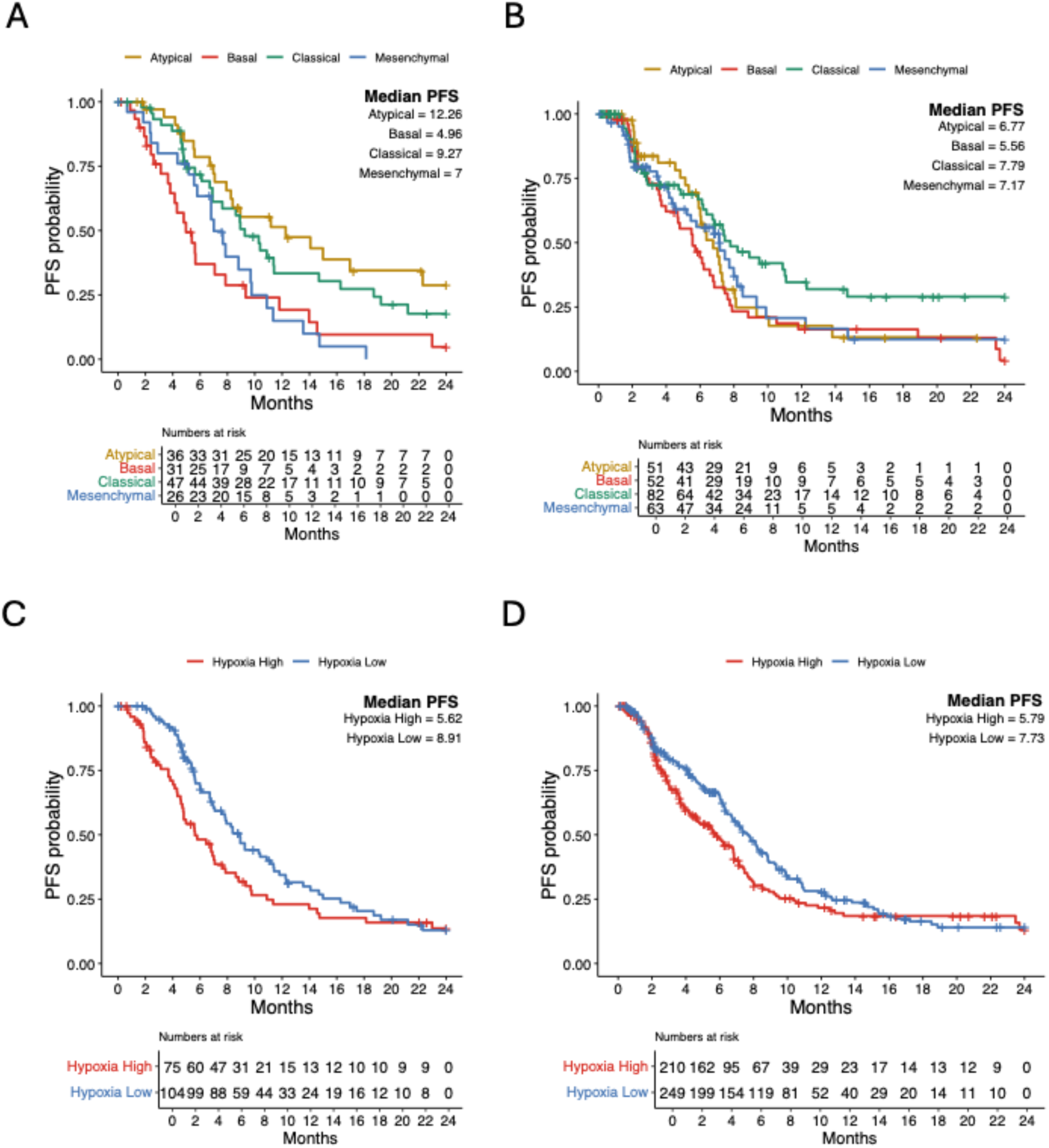
Treatment-specific progression-free survival (PFS) patterns in traditional and hypoxia subtypes across clinical settings. PFS curves for patients grouped by traditional subtypes in the **(A)** 1L setting and **(B)** M1L setting. PFS curves for patients grouped by hypoxia subtypes in the **(A)** 1L setting and **(B)** M1L setting.

**Supplementary Figure S9.**
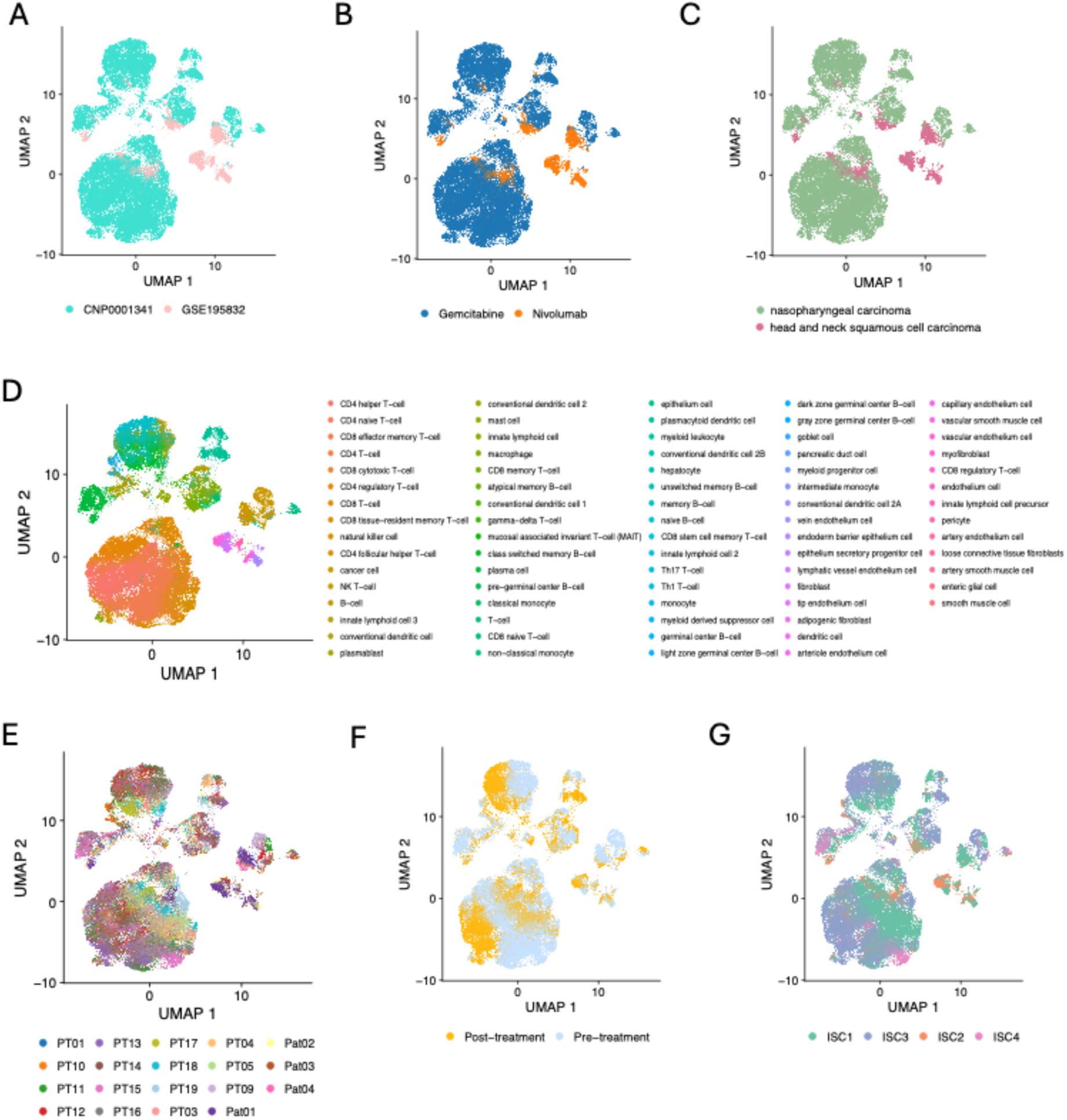
Single-cell landscape of two studies featuring pre– and post-treatment samples from HNSCC patients treated with Chemo and CPI. **(A)** UMAP plot showing integrated single-cell RNA-seq data from two publicly available datasets (CNP0001341 and GSE195832), demonstrating successful cross-study integration. **(B)** Cells colored by treatment regimen, distinguishing patients treated with **Gemcitabine** (Chemo) versus **Nivolumab** (CPI). **(C)** Cells grouped by cancer type. **(D)** Cell type annotations based on Nebion reference. **(E)** UMAP colored by individual patient IDs. **(F)** Treatment status– specific distribution of cells. **(G)** Immune subtype classification (ISC1–ISC4) mapped onto the integrated single-cell space.

**Supplementary Figure S10:**
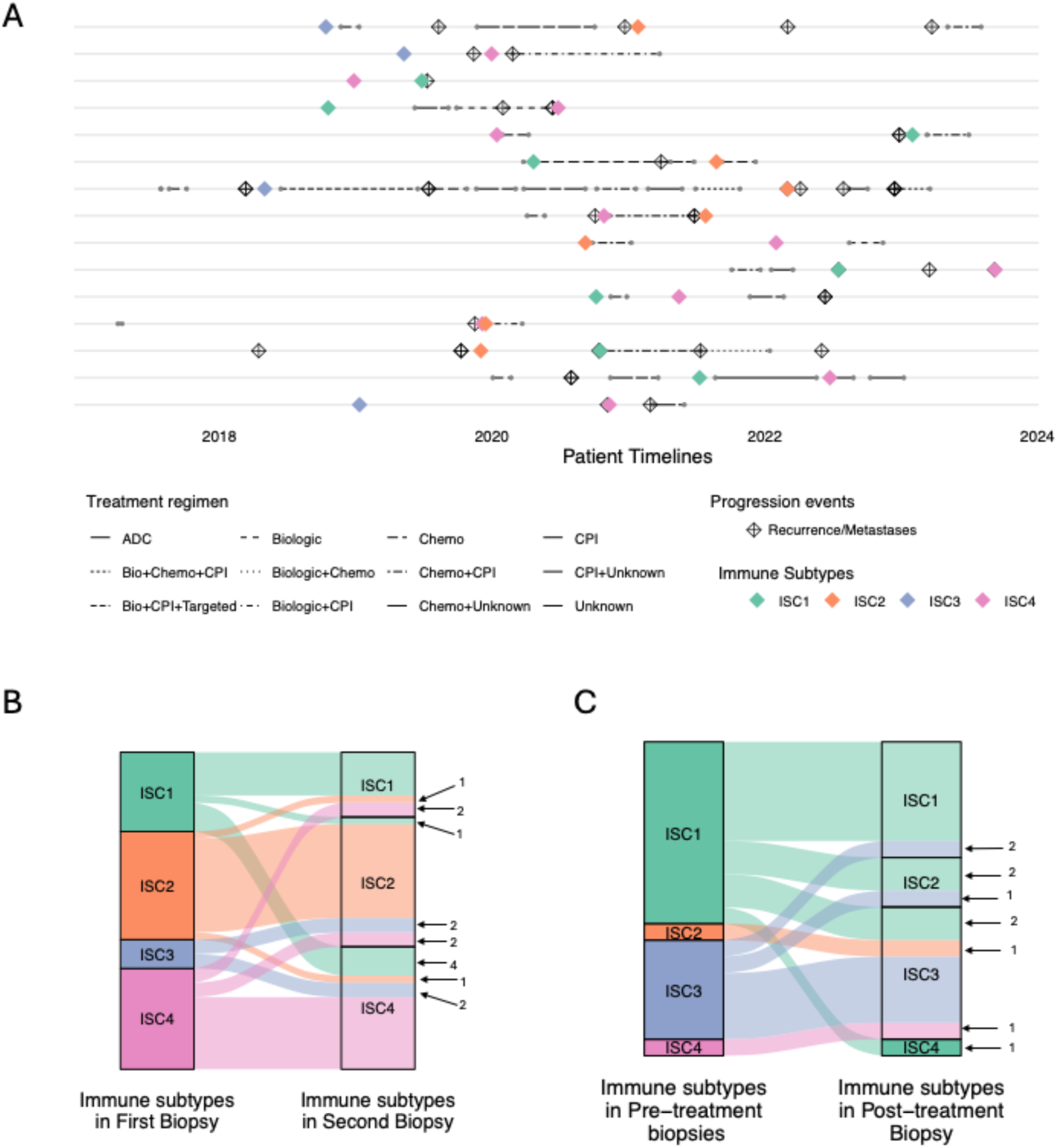
Immune Subtype Dynamics Across Timepoints and Modalities. **(A)** Patient-level timelines showing treatment history, recurrence/metastasis events, and immune subtypes at biopsy timepoints. Each row represent a single patient, with lines indicating treatment regimens (line types) over time. **(B)** Sankey diagram illustrating changes in immune subtypes between the first and second biopsies in the real-world evidence (RWE) cohort. **(C)** Sankey diagram showing immune subtype transitions between pre-treatment and post-treatment biopsies in the single-cell cohort. Frequencies of transitions are shown.

**Supplementary Figure S11:**
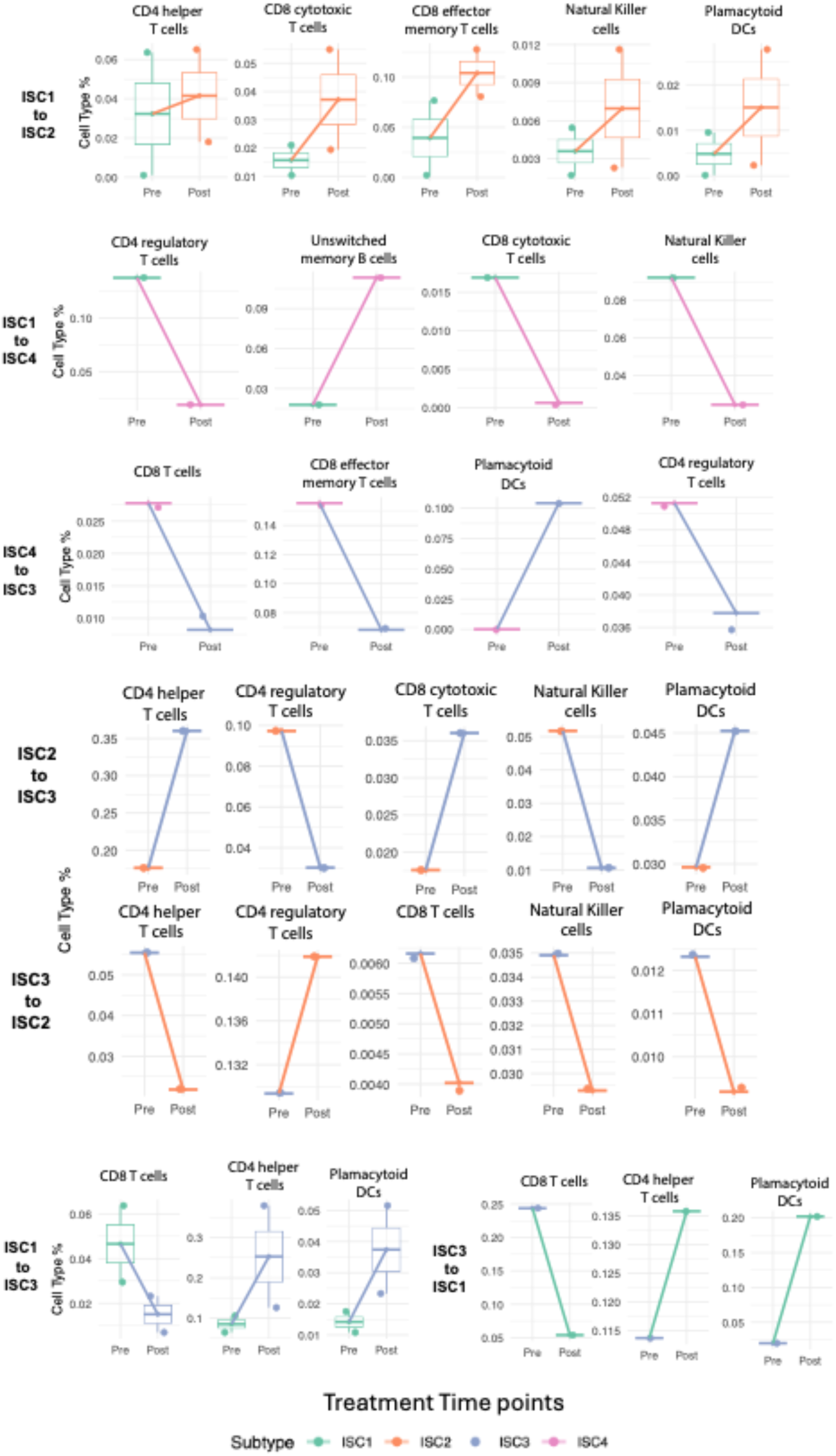
Immune cell-type remodeling across treatment-associated subtype transitions in HNSCC. Boxplots and paired lines represent changes in the relative abundance of selected immune cell types between pre– and post-treatment biopsies, stratified by immune subtype transition. Each facet corresponds to a unique subtype–cell type pair (e.g., ISC1→ISC2: CD8 effector memory T cells). Lines connect pre– and post-treatment samples per patient, colored by final (post-treatment) subtype. Cell-type frequencies were calculated as a percentage of total cells per patient and treatment time point using single-cell data from HNSCC tumors. Subtype transitions were inferred by comparing subtype labels between pre– and post-treatment samples for each patient. Cell types shown were selected based on the largest absolute changes in relative abundance (|mean change| > 0.05) across transition groups. Only transitions and cell types meeting this threshold were retained for visualization.

